# Deep learning enables rapid and robust analysis of fluorescence lifetime imaging in photon-starved conditions

**DOI:** 10.1101/2020.12.02.408195

**Authors:** Yuan-I Chen, Yin-Jui Chang, Shih-Chu Liao, Trung Duc Nguyen, Jianchen Yang, Yu-An Kuo, Soonwoo Hong, Yen-Liang Liu, H. Grady Rylander, Samantha R. Santacruz, Thomas E. Yankeelov, Hsin-Chih Yeh

**Affiliations:** Department of Biomedical Engineering, The University of Texas at Austin, Austin, TX 78712, USA; ISS, Inc., 1602 Newton Drive, Champaign, IL 61822, USA; Texas Materials Institute, The University of Texas at Austin, Austin, TX 78712, USA; Institute for Neuroscience, The University of Texas at Austin, Austin, TX 78712, USA; Oden Institute for Computational Engineering and Sciences, The University of Texas at Austin, Austin, TX 78712, USA; Department of Diagnostic Medicine, The University of Texas at Austin, Austin, TX 78712, USA; Department of Oncology, The University of Texas at Austin, Austin, TX 78712, USA; Livestrong Cancer Institutes, The University of Texas at Austin, Austin, TX 78712, USA; Master Program for Biomedical Engineering, China Medical University, Taichung 40402, TAIWAN; Research Center for Cancer Biology, China Medical University, Taichung 40402, TAIWAN

## Abstract

Fluorescence lifetime imaging microscopy (FLIM) is a powerful tool to quantify molecular compositions and study the molecular states in the complex cellular environment as the lifetime readings are not biased by the fluorophore concentration or the excitation power. However, the current methods to generate FLIM images are either computationally intensive or unreliable when the number of photons acquired at each pixel is low. Here we introduce a new deep learning-based method termed *flimGANE* (fluorescence lifetime <u>im</u>aging based on <u>G</u>enerative <u>A</u>dversarial <u>N</u>etwork <u>E</u>stimation) that can rapidly generate accurate and high-quality FLIM images even in the photon-starved conditions. We demonstrated our model is not only 258 times faster than the most popular time-domain least-square estimation (*TD_LSE*) method but also provide more accurate analysis in barcode identification, cellular structure visualization, Förster resonance energy transfer characterization, and metabolic state analysis. With its advantages in speed and reliability, *flimGANE* is particularly useful in fundamental biological research and clinical applications, where ultrafast analysis is critical.

## Introduction

Using fluorescence decay rate as the contrast mechanism, fluorescence lifetime imaging microscopy (FLIM) is a powerful quantitative tool for studying cell and tissue biology^1, 2, 3^, allowing us to monitor the pH^4^, viscosity^5^, temperature^6^, oxygen content^7^, metabolic state^8^ and functional property of a biomarker^9^ inside live cells or tissues. Depending on the intrinsic property of fluorophore, FLIM images are not skewed by fluorophore concentration and excitation power, eliminating the biases introduced by the traditional intensity-based images^2^. Combined with the fluorescence resonance energy transfer (FRET) sensors^10^, FLIM can probe Ca^2+^ concentration^11^, glucose concentration^12^ and protein-protein interactions^13^, without the need to measure acceptor’s fluorescence^14^. Whereas FLIM offers many unique advantages in quantifying molecular interactions^15^ and chemical environments^16^ in biological or chemical samples, fluorescence lifetime analysis is a slow process with results often impaired by fitting errors. Adopted from disparate disciplines, various fluorescence lifetime estimation methods such as curve fitting (least-squares fitting^17^, maximum likelihood estimation^18^, global analysis^19^ and Bayesian analysis^20^), phasor analysis^21, 22^ and deconvolution analysis (stretched exponential analysis^23^, Laguerre deconvolution^24^) have been developed to infer the lifetime of interest. However, different methods are limited by poor accuracy particularly in low-light conditions, long computation times or susceptible to error from initial assumption of decay parameters.

Here we demonstrate a new fluorescence lifetime <u>im</u>aging method based on <u>G</u>enerative <u>A</u>dversarial <u>N</u>etwork <u>E</u>stimation (*flimGANE*) that can provide fast, fit-free, precise, and high-quality FLIM images even under the extreme low-light conditions. GAN is one of the frameworks for evaluating generative models *via* an adversarial process^25^, which have been adopted to improve astronomical images^26, 27^, transform images across different modalities^26, 28^, and design drugs that target specific signaling molecules^29^. While GAN-based algorithms have recently drawn much attention for inferring photo-realistic natural images^30^, they have not been used to generate high-quality FLIM images based on the fluorescence decays collected by a laser scanning confocal microscope. Our *flimGANE* method is adapted from the Wasserstein GAN algorithm^31^ (WGAN; see Methods), where the generator (*G*) is trained to produce an “artificial” high-photon-count fluorescence decay histogram based on a low-photon-count input, while the discriminator (*D)* distinguishes the artificial decay histogram from the ground truth (which can be a simulated dataset or a decay histogram collected under strong excitation). As a minimax two-player game, the training procedure for *G* is to maximize the probability of *D* making a mistake^25^, eventually leading to the production of very realistic, artificial high-photon-count decay histograms that can be used to generate a high-quality FLIM image. Using a well-trained generator (*G*) and an estimator (*E*), we can reliably map a low-quality decay histogram to a high-quality counterpart, and eventually to the three lifetime parameters (*α*_*1*_, *τ*_*1*_, and *τ*_*2*_) within 0.32 ms/pixel (see Methods). Without the need to do any curve fitting based on initial guesses, our *flimGANE* method is 258-fold faster than the time-domain least-squares estimation (*TD_LSE*^*32, 33*^) and 2,800-fold faster than the time-domain maximum likelihood estimation (*TD_MLE*^34, 35^) in generating a 512 512 FLIM image. While almost all commercial FLIM analysis tools are based on *TD_LSE*, using the least-squares estimator to analyze Poisson-distributed data is known to lead to biases^36^, making *TD_MLE* the gold standard for FLIM analysis by many researchers^18^. Our *flimGANE* can provide similar FLIM image quality as *TD_MLE*, but much faster.

Overcoming a number of hardware limitations in the classical analog frequency domain approach, the digital frequency-domain (DFD) lifetime measurement method has substantially increased the FLIM analysis speed^21, 22, 37^. The acquired DFD data at each pixel, termed a cross-correlation phase histogram, can lead to a phasor plot with multiple harmonic frequencies. From such a phasor plot, modulation ratio and phase shift at each harmonic frequency can be obtained, which are then fitted with a least-squares estimator (*LSE*) to generate a lifetime at each pixel (termed the *DFD_LSE* method). Our *flimGANE* not only runs nearly 12-fold faster than *DFD_LSE* but also produces more accurate quantitative results and sharper structural images of *Convallaria* and live HeLa cells. Whereas the lowest number of photons needed for reliable estimation of a fluorescence lifetime by *TD_MLE* is about 100 photons^38^, *flimGANE* performs consistently well with a photon count as low as 50 per pixel in our simulations. Moreover, *flimGANE* improves the energy transfer efficiency estimate of a glucose FRET sensor, leading to a more accurate glucose concentration measurement in live HeLa cells. Providing both efficiency and reliability in analyzing low-photon-count decays, our *flimGANE* method represents an important step forward towards real-time FLIM.

## Results

### Training the generative adversarial network in *flimGANE*

Based on the Wasserstein GAN framework (see Methods), *flimGANE* is designed to analyze one- or two-component fluorescence decays under photon-starved conditions (**Fig. 1**). There are two ways to generate a dataset of ground-truth lifetime histograms for training *G* and *D* – either by creating a decay dataset using Monte Carlo (MC) simulations or by acquiring an experimental dataset from standard organic fluorophores under high excitation power. The inputs of *G* are degraded data from the ground truths, which can be obtained by running simulations at a low-emission rate or by recollecting experimental data under low excitation power.

**Fig. 1.**
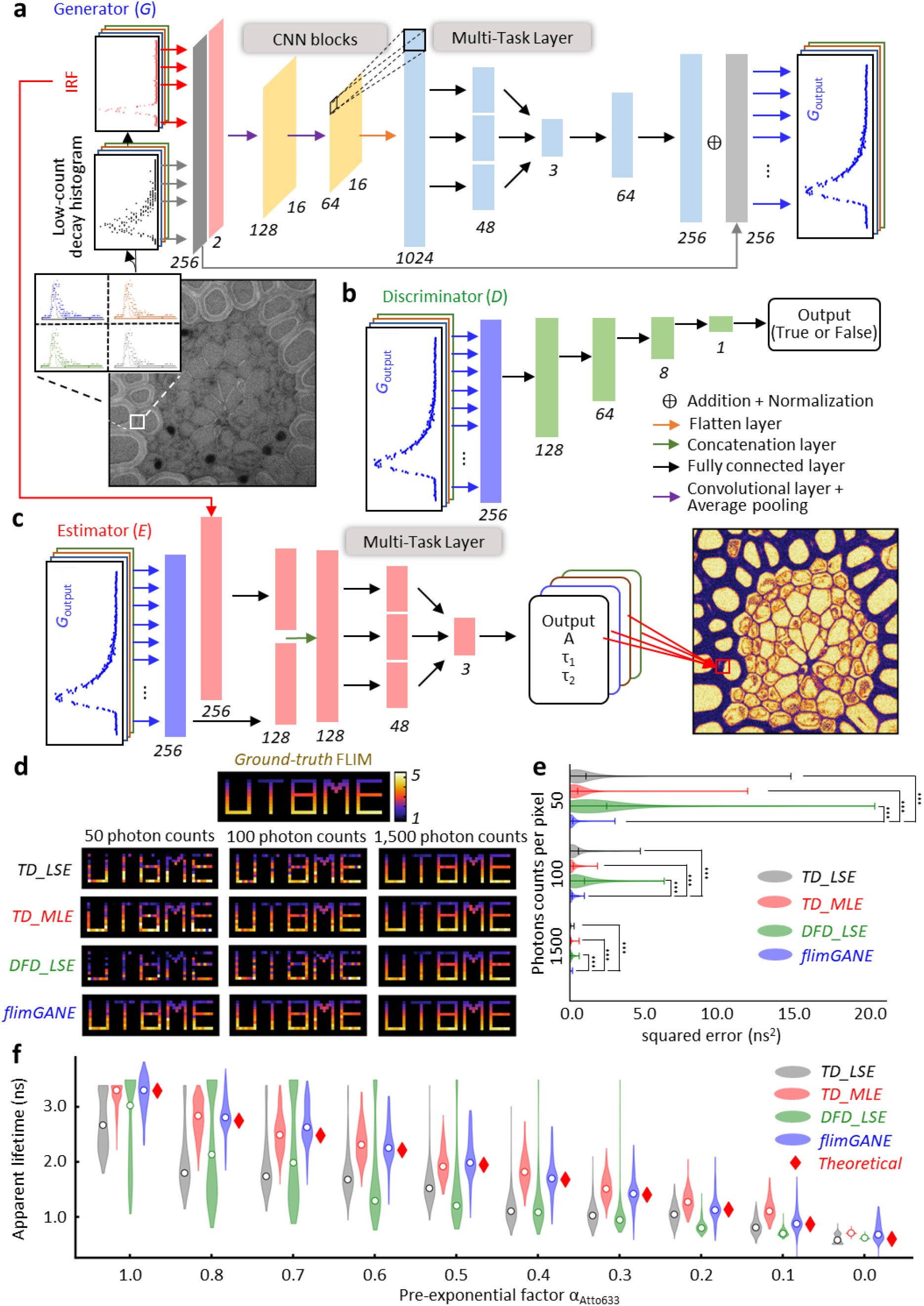
*flimGANE* (fluorescence lifetime<u>im</u>aging based on<u>G</u>enerative<u>A</u>dversarial<u>N</u>etwork<u>E</u>stimation) is a rapid and accurate method to generate fluorescence lifetime microscopy (FLIM) images. (**a-c**) Schematic of deep learning framework for *flimGANE* architecture. (**a**) The generator section is used to transform the acquired decay curve into a higher-count one. It comprises two CNN blocks, each of which are made up of one convolutional layer followed by an average pooling layer of stride two. The CNN section is followed by a flatten layer. Then a multi-task layer converts data into virtual lifetime parameters, followed by two fully-connected layers. Skip connection is used to pass data between layers of the same level. (**b**) The discriminator consists of four fully connected layers. (**c**) The estimator comprises a partially connected and a fully connected layer followed by a multi-task layer to map the high-count decay curve into lifetime parameters. (**d**) Comparison of FLIM image generated by different methods (n = 134 pixels). (**e**) Squared error with the ground truth comparison of different methods. Under extremely low photon count condition (50 counts), only *flimGANE* successfully provides accurate estimation. Under low photon count condition (100 counts), *TD_LSE* and *DFD_LSE* failed to generate accurate FLIM image; while under high photon count condition (1500 counts), all the FLIM images match well with the ground truth. Significant differences are indicated as: *** (p < 0.001). (**f**) Our model successfully characterizes apparent lifetime of the mixture of two fluorescent dyes (stock solution: 3 μM Cy5-NHS ester and 7 μM Atto633 in DI water) with 10 different ratios (20*20 pixels). The mean values obtained from Gaussian fitting are indicated as white solid circles.

We started our network training using an MC simulation dataset (**Supplementary Fig. 1**). A Python program was employed to simulate the photon collection process in the counting device with 256 time bins, following the probability mass function (pmf) numerically calculated by the convolution of an experimentally obtained instrument response function (IRF) and a theoretical two-component decay model (*α*_*1*_, *τ*_*1*_, 1-*α*_*1*_ and *τ*_*2*_) at a selected emission rate (*rate*)^39^. Depending on the fluorophores that users want to image, proper *α*_*1*_, *τ*_*1*_, *τ*_*2*_ and *rate* parameters that span the range of interest could be selected (**Supplementary Table 1**), generating about 600 normalized ground truths and 300k degraded decays for training *G* and *D*. The adversarial network training was completed in 6.1 hours (see Methods**; Figs. 1a-b**; **Supplementary Fig. 2**). The normalized degraded decay was transformed into the normalized “ground-truth mimicking” histogram, termed *G*_*output*_ (**Supplementary Fig. 3**), within 0.17 ms. Such a *G*_*output*_ was indistinguishable from the ground truths by *D. E*, which was separately trained on the ground truths and completed in 0.1 hours, was then employed to extract the key lifetime parameters (*α*_*1*_, *τ*_*1*_, and *τ*_*2*_) from the *G*_*output*_ within 0.15 ms (**Fig. 1c**). Then, the combining training of the *G* and *E* took extra 0.7 hours, in order to adapt the pre-trained *E* to the current *G*_*output*_ (**Supplementary Fig. 2**).

To demonstrate the reliability of our *flimGANE* method, we created a set of 14×47 “*UTBME*” FLIM images *in silico* (independently generated, not used in the training process) at three photon emission rates (50, 100 and 1,500 photons per pixel). At 1,500 photons per pixel, all four methods (*TD_LSE, TD_MLE, DFD_LSE* and *flimGANE*) generated high-fidelity FLIM images (based on the apparent lifetime, *τ*_*α*_ = *α*_*1*_*τ*_*1*_ + (1-*α*_*1*_)*τ*_*2*_, see Methods), with mean-squared errors (MSE) less than 0.10 ns^2^. At 100 photons per pixel, *flimGANE* had similar performance as *TD_MLE* (MSE were both less than 0.20 ns^2^); however, *flimGANE* clearly outperformed *TD_LSE, TD_MLE*, and *DFD_LSE* at 50 photons per pixel (0.19 vs. 1.04, 0.49 and 2.41 ns^2^, respectively; **Fig. 1d-e, Supplementary Figs. 4**-**6**, and **Supplementary Tables 2**-**5**). Speed analysis showed that *flimGANE* was 258 and 2,800 times faster than *TD_LSE* and *TD_MLE*, respectively (*flimGANE* – 0.32 ms per pixel, *TD_LSE* – 82.40 ms, *TD_MLE* – 906.37 ms; **Supplementary Table 6**). While *DFD_LSE* offered a relatively high speed in generating FLIM images (3.94 ms per pixel), its accuracy was worse than that of *flimGANE* (**Figs. 2**-**5**). In contrast, being a computationally intensive method, *TD_MLE* offered the accuracy, but not the speed. Only *flimGANE* could provide both speed and accuracy in generating FLIM images. In addition, the MLE method became unreliable in the extremely low-photon-count condition (50 photons per pixel), while *flimGANE* still provided a reasonable result.

To obtain accurate FLIM images, the IRF of the imaging system, which is mainly decided by the width of laser pulse and the timing dispersion of detector, should be carefully considered during lifetime estimation. While the FWHM of IRF is stable in most of the commercial FLIM imaging systems (detector time jittering within 35-500 ps^40^), users often observe that the delay between the single-photon detector output and the TCSPC electronics input varies from day to day, possibly due to the instability of the TCSPC electronics caused by radio-frequency interference, laser lock instability, and temperature fluctuation. Such delay changes cause the onsets of the decays to drift, deteriorating the *flimGANE* analysis results. A preprocessing step, termed Center of Mass Evaluation (CoME), was thus introduced to adjust (or standardize) the temporal location of the onset of experimental decays (**Supplementary Figs. 7**-**9**). After preprocessing, the apparent lifetimes estimated by *flimGANE* were found free of onset-delay bias.

To prove the reliability of *flimGANE* in estimating an apparent fluorescence lifetime from a mixture, two fluorophores, Cy5-NHS ester (*τ*_*1*_ = 0.60 ns) and Atto633 (*τ*_*2*_ = 3.30 ns), were mixed at different ratios, creating ten samples of distinct apparent fluorescence lifetimes (*τ*_*α*_) ranging from 0.60 to 3.30 ns. Here *τ*_*1*_ and *τ*_*2*_ were measured from the pure dye solutions and estimated by *TD_MLE*, whereas the theoretical apparent lifetime *τ*_*αT*_ was predicted by the equation *τ*_*αT*_ = *τ*_*1*_*α*_*1*_ + *τ*_*2*_(1-*α*_*1*_). *α*_*1*_, the pre-exponential factor^41^, was derived from the relative brightness of the two dyes and their molar ratio ^37^ (see Methods). When analyzing 256×256-pixel images with emission rates fluctuating between 80-200 photons per pixel, *flimGANE* and *TD_MLE* produced the most accurate and precise *τ*_*α*_ estimates among the 4 methods (**Fig. 1f**, and **Supplementary Table 7**). *TD_LSE* and *DFD_LSE* performed poorly in this low-light, two-dye mixture experiment.

### Discriminating fluorescence lifetime barcode beads

We then tested *flimGANE* in discriminating the fluorescence lifetime barcodes. To create fluorescence lifetime barcodes, biotinylated Cy5- and Atto633-labeled DNA probes were mixed at three different ratios, Cy5-DNA:Atto633-DNA = 1:0 (*barcode_1*, expecting lifetime 1.90 ns); 1:1 (*barcode_2*, 2.40 ns) and 0:1 (*barcode_3*, 3.50 ns), and separately conjugated to streptavidin-coated polystyrene beads (3-4 μm in size, see **Methods**). It was noted that the lifetime of Cy5-DNA (1.90 ns) is different from that Cy5-NHS ester (0.60 ns). Similarly, the lifetime of Atto633-DNA (3.50 ns) is different from that of Atto633 (3.30 ns). A cover slip coated with the three barcode beads (at equal molar concentration) was scanned by the ISS Alba v5 confocal microscopic system (equipped with a 20 MHz 635 nm diode laser for excitation and a *FastFLIM* module for DFD acquisition^37^) for 31 seconds, generating 512×512-pixel DFD data with photon counts ranging from 50-300 per pixel on the beads (**Fig. 2a**). The acquired DFD data (i.e., cross-correlation phase histograms^37^) were converted into time decays for *flimGANE, TD_LSE* and *TD_MLE* analysis (**Fig. 2b**). Each barcode bead was registered by ImageJ ROI manager and assigned an ID number (**Supplementary Fig. 10**). An apparent lifetime was assigned to each pixel on the bead (∼292 pixels) and lifetimes of all pixels were plotted in a histogram. The mean lifetime for the bead was determined by the Gaussian fit of the histogram. After examining 97 beads, we chose the cutoff lifetimes to be 2.15 and 2.95 ns for barcode identification (**Fig. 2c**) and assigned pseudocolors to the barcode beads (**Fig. 2d**). It was clear to see that *flimGANE* is the only method that can correctly identify the three barcodes and restore the 1:1:1 barcode ratio, while other methods often misidentified the barcodes (**Fig. 2e**). Whereas it was a general trend that beads with more Atto633-DNA are dimmer, possibly due to stronger self-quenching, brightness alone could not classify the three barcodes (**Fig. 2c, Supplementary Fig. 11, Supplementary Table 8**). It was noted that the brightness of *barcode_1* beads could vary by six-fold, but the coefficient of variance (CV) in *barcode_1* lifetimes was only 0.06, making lifetime a better metric to differentiate barcodes.

**Fig. 2.**
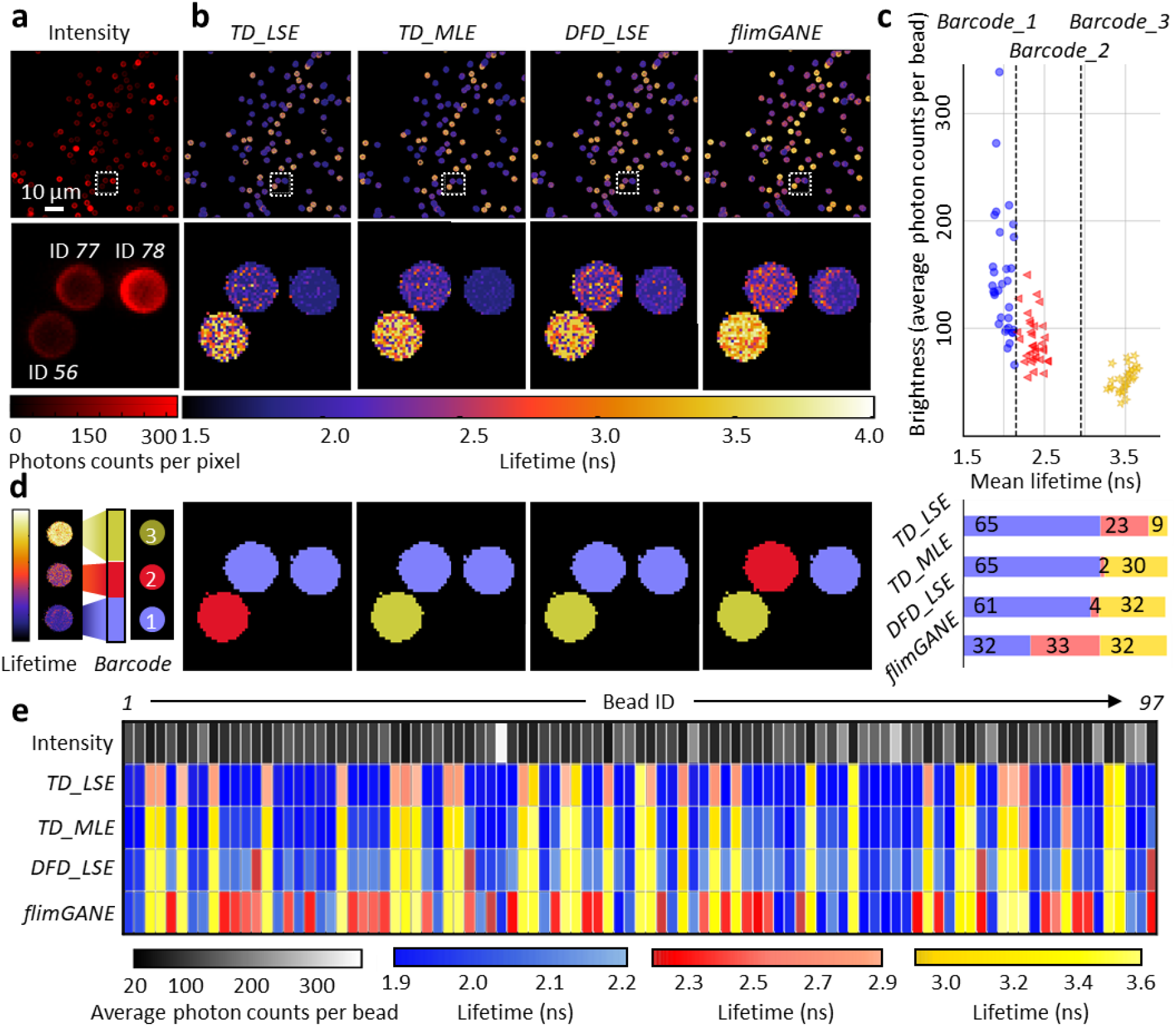
*flimGANE* accurately classifies the three fluorescence barcodes. (**a**) A contrast image shows 3-µm fluorescent beads with various brightness, where (**b**) FLIM images obtained from *TD_LSE, TD_MLE, DFD_LSE, flimGANE* reveal three lifetime species. (**c**) Mean lifetimes vs. brightness of 97 beads shows the intensity differences of the 3 barcodes and the cutoff lifetimes (2.15 and 2.95 ns) for barcode identification. (**d**) Barcode classification results by the four methods indicate that only *flimGANE* can correctly identify the barcodes and restore the correct barcode ratio (1:1:1). (**e**) Classification results of all 97 beads show that many *barcode_2* beads were misidentified as *barcode_1* beads by *TD_LSE, TD_MLE* and *DFD_LSE*, while the identification of *barcode_3* is mostly reliable among these methods (except for *TD_LSE*, which performs poorly under the low-light conditions).

### Visualizing cellular structures of *Convallaria* and HeLa cells

The DFD fluorescence data of *Convallaria* (lily of the valley) and live HeLa cells, acquired under the low- and the medium-photon-count conditions (**Fig. 3a**), were analyzed by *DFD_LSE* and *flimGANE* (**Fig. 3b**), where *TD_MLE* (in medium-photon-count condition, ∼243 photons per pixel) served as the standard for comparison. The histogram clearly showed two characteristic lifetimes (0.90 ± 0.13 ns; 4.84 ± 1.20 ns) in the *Convallaria* sample (**Fig. 3c**). As *TD_MLE* with medium photon counts had all lifetime estimates within 0.0-6.0 ns range, we limited the upper bound of the lifetime estimates in to be 6.0 ns. Those 6.0 ns pixels were given the white pseudocolor and regarded as failed pixels in the FLIM images (**Fig. 3d**). A large number of failed pixels were seen in the *DFD_LSE* images (37% and 25% for the low- and medium-count images, respectively; **Fig. 3b**), deteriorating the visualization of structure details in the *Convallaria* sample. In contrast, there were very few failed pixels in the *flimGANE* images under the low-light condition (∼83 photons per pixel), making them most resemble the *TD_MLE* images under medium-light condition and provide better visualization of the structure details (**Figs. 3c-d**). The structure similarity index (SSIM)^42^ indicated that the *flimGANE* images were 73% more similar to the gold standard *TD_MLE* images than those generated by *DFD_LSE* (*flimGANE* – 0.88, *DFD_LSE* – 0.51; **Supplementary Table 9**), and visual information fidelity (VIF)^43^ showed that the *flimGANE* images were 1.44-fold higher than those reconstructed by *DFD_LSE* (*flimGANE* – 0.22, *DFD_LSE* – 0.09; **Supplementary Table 9**).

**Fig. 3.**
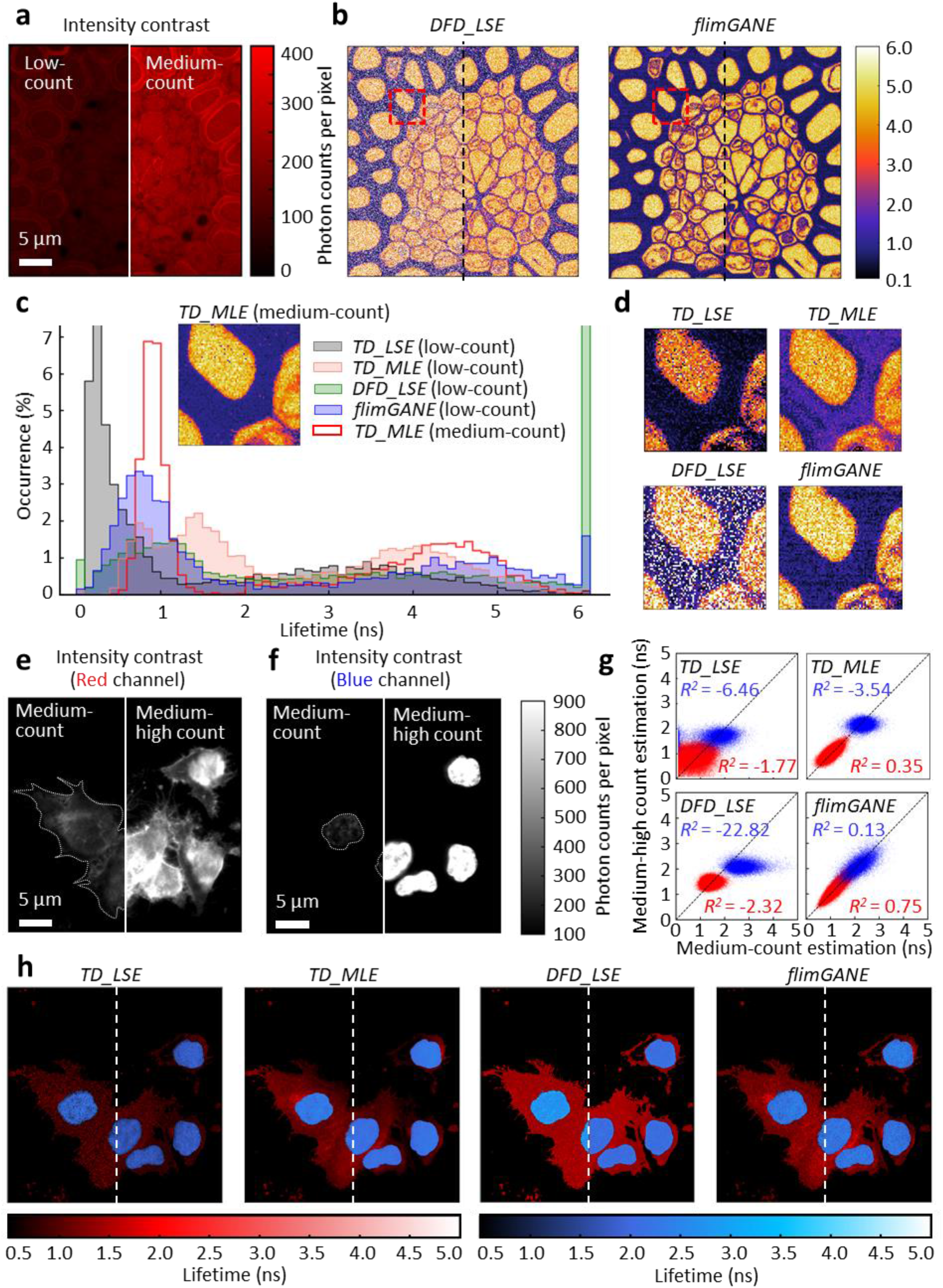
*flimGANE* provides better and more consistent visualization of *Convallaria* and HeLa cell samples. (**a**) Intensity contrast of *Convallaria* was imaged with a size of 512 x 512 pixels and an intensity ranged from 50 – 150 counts (left; low photon counts) and from 300 – 400 counts (right; medium photon counts). (**b**) FLIM images generated by *DFD_LSE* and by *flimGANE* demonstrated that *flimGANE* was more robust than *DFD_LSE*, where the estimation was independent of photon counts. (**c**) Histogram of lifetime obtained from *TD_LSE, TD_MLE, DFD_LSE, flimGANE*, and medium-count based *TD_MLE* in the selected ROI showed that *flimGANE* showed the most similar distribution with the standard. (**d**) A zoomed-in ROI (red box; low photon counts) was selected and analyzed with *TD_LSE, TD_MLE, DFD_LSE* and *flimGANE* to reveal further details of the structure. (**e**) Intensity contrast of plasma membrane of live HeLa cells was imaged with a size of 512 x 512 pixels in red channel (685/40 nm,) and an intensity ranged from 80 – 400 counts (left; medium photon counts) and from 300 – 1,000 counts (right; medium-high photon counts). Dash line represents the contour of live cells. (**f**) Intensity contrast of nuclei of live HeLa cells was imaged with a size of 512 x 512 pixels in blue channel (494/34 nm) and an intensity ranged from 50 – 300 counts (left; medium photon counts) and from 300 – 1,500 counts (right; medium-high photon counts). (**g**) 2D scatter plots of lifetime acquired at low and medium excitation power. *flimGANE* provided more consistent estimates at two different photon rates. The coefficient of determination, *R*^*2*^, ranged from −◸ to 1.00, was utilized to evaluate the consistency by setting the true value of medium-high-count estimations as the medium-count ones. (**h**) Overlay of FLIM images in red and blue channels (left; medium photon counts; right; medium-high photon counts).

In the live HeLa cell sample, nuclei and membranes were stained with Hoechst and CellMask™ Red and excited by 405 nm and 635 nm diode lasers, respectively. The contours of nuclei and cell membranes could not be clearly defined by the intensity-based images even under medium-light condition (**Figs. 3e-f**). Although FLIM overlay images allowed us to better visualize structural details in HeLa cells, the lifetime estimates could be biased even when there were ∼180 photons per pixel (medium-light condition; *TD_LSE* and *DFD_LSE* images in **Fig. 3h**). Using the medium-high-count *TD_MLE* images (∼600 photons per pixel) as the standard for comparison, *flimGANE* clearly outperformed *TD_LSE, TD_MLE*, and *DFD_LSE* in producing images that resemble the standard under medium-light condition (**Fig. 3h; Supplementary Table 10**). Interestingly, when scrutinizing the assigned lifetime at each pixel, we found not only *TD_LSE* and *DFD_LSE* but also *TD_MLE* give inconsistent lifetime estimates at the two excitation powers (e.g., *R*^*2*^ in blue channel were −6.46, −3.54 and −22.82 for *TD_LSE, TD_MLE* and *DFD_LSE*, respectively). In contrast, *flimGANE* provided much more consistent lifetime estimates regardless the excitation power (*R*^*2*^ was 0.13 in blue channel; **Fig. 3g**).

### Quantifying Förster resonance energy transfer (FRET) efficiency in live MDA-MB-231 cells

Combined with the glucose FRET sensor, FLIM has been employed to image the glucose concentration in live cells^10, 44^. However, depending on the lifetime analysis methods, the trend of FRET change can be skewed, especially when the donor lifetime change is very small (e.g., only 0.1-0.2 ns). Our glucose FRET sensor, termed CFP-*g*-YFP^45^, consisted of a glucose binding domain flanked by a cyan fluorescent protein (CFP) donor and a yellow fluorescent protein (YFP) acceptor (see Methods, **Fig. 4a**). The overlap between CFP emission and YFP absorption leads to efficient dipole-dipole interactions. The CFP-*g*-YFP sensor-expressed MDA-MB-231 tumor cells were starved for 24 hrs before adding different amount of glucose to the cell culture (final concentrations: 0, 0.5, 1.0, 2.0, 5.0, 10.0, 15.0 mM). The confocal scanning system collected DFD data from a 256×256-pixel area before and after the addition of glucose, which were then analyzed by *TD_LSE, TD_MLE, DFD_LSE*, and *flimGANE* methods to generate FLIM images based on the CFP donor decays (**Fig. 4b**). By proper selection of regions of interest (ROI) in imaging analysis, single cells were separated from each other and from the background noise (**Supplementary Fig. 12; Supplementary Table 11**). Thousands of lifetime data points (apparent lifetimes, *τ*_*a*_) were plotted in a histogram and the mean was extracted by Gaussian fitting, giving one representative donor lifetime for each glucose concentration (**Fig. 4c**). The energy transfer efficiency (*E*) was calculated based on the equation: *E = 1-(τ*_*DA*_ */τ*_*D*_), where *τ*_*D*_ and *τ*_*DA*_ were the representative CFP lifetimes before and after addition of glucose, respectively. Whereas only subtle lifetime changes were seen in the CFP donor lifetime (0.04-0.20 ns in **Supplementary Table 12**, which led to low FRET efficiencies around 0.02-0.07), *flimGANE*-derived FRET efficiencies were not only highly reproducible but also showing a general increasing trend at higher glucose concentrations. On the other hand, the lifetime of acceptor (YFP) did not change upon addition of glucose (**Supplementary Fig. 13**). Among the four methods, *DFD_LSE* failed to provide a FRET efficiency response curve due to its poor lifetime estimation in this experiment, thus being excluded from **Fig. 4c**.

**Fig. 4.**
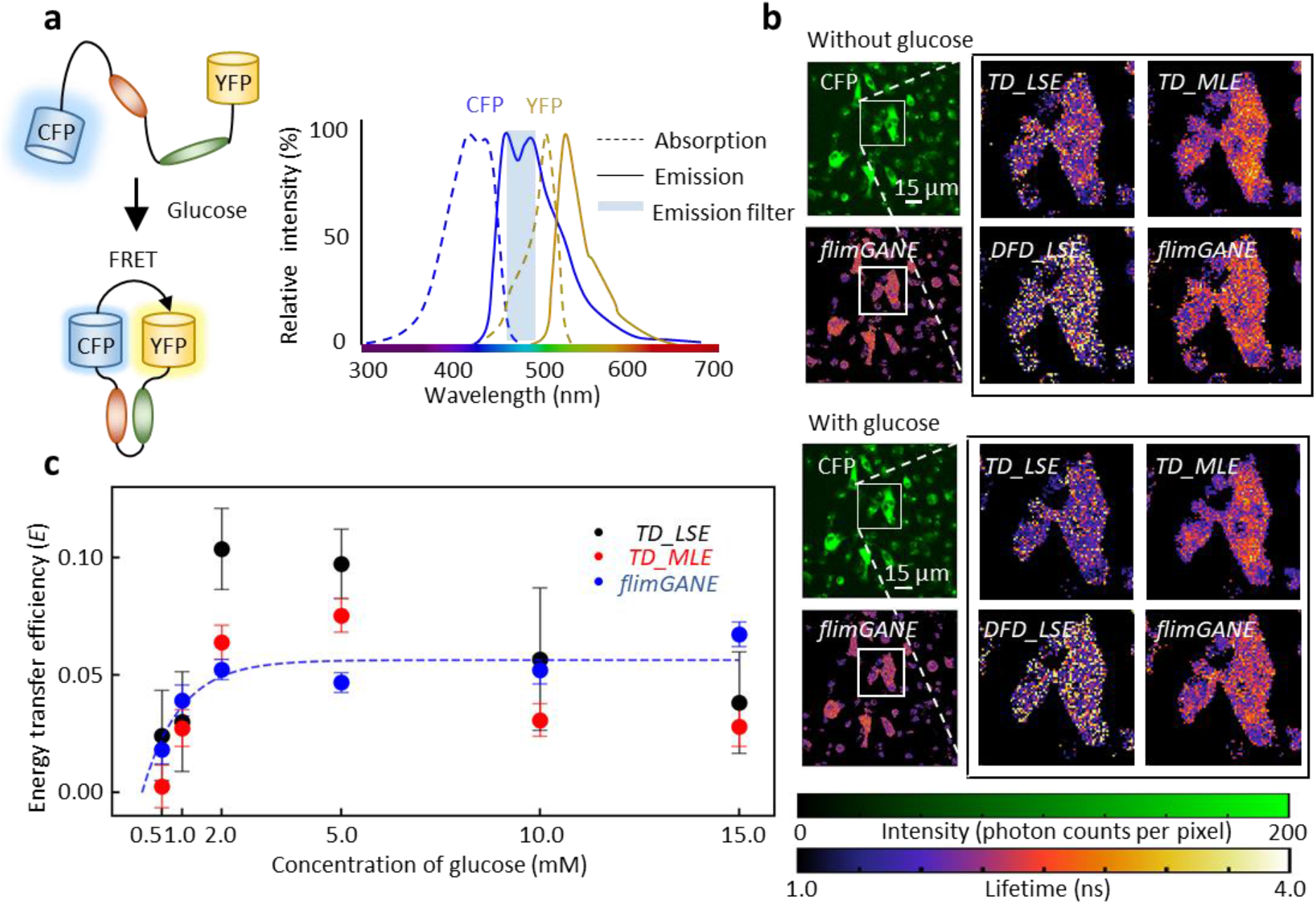
Deep-learning enabled FRET measurement from low-photon-count FLIM imaging. (**a**) Normalized excitation and emission spectra of CFP and YFP. Dotted rectangles indicate transmission of emission filters and schematic of CFP-*g*-YFP FRET pair interaction with glucose. (**b**) Intensity contrast, FLIM images generated by *TD_LSE, TD_MLE, DFD_LSE* and *flimGANE* of CFP are presented before adding 2 mM glucose, and immediately after adding 2 mM glucose. (**c**) Energy transfer efficiency, *E*, was plotted the versus the concentration of Glucose that was added (error bars, standard deviation errors on the parameter estimate, n = 1507 ∼ 6824). An asymptotic phase of sigmoidal curve fitted well with the observation from *flimGANE* (*R*^*2*^ = 0.92).

While the intensity-based method, *E = 1-(F*_*DA*_ */F*_*D*_), was used to estimate *E*, the resulting response curve clearly deviated from the reasonable trend, possibly due to the artifacts such as photobleaching. When comparing the CFP FLIM images at 2 mM glucose concentration, we could clearly see that the *flimGANE* image well resembled the *TD_MLE* image, but not the other two images, in which there were many failed pixels (**Fig. 4b**). Although the *TD_MLE* images were similar to the *flimGANE* images, *TD_MLE*-derived FRET efficiencies had higher variations and showed an unrealistic, decreasing trend at higher glucose concentrations. In this demonstration, *flimGANE* not only gave a correct sensor response curve but also provided an analysis speed 2,800-fold faster than *TD_MLE* in reconstructing a FLIM image.

### Quantifying metabolic states in live HeLa cells

Autofluorescence of endogenous fluorophores, such as nicotinamide adenine dinucleotide (NADH), nicotinamide adenine dinucleotide phosphate (NADPH), and flavin adenine dinucleotide (FAD), are often used to characterize the metabolic states of individual cancer cells, through metrics such as optical redox ratio (ORR)^46^, optical metabolic imaging index (OMI index)^47^ and fluorescence lifetime redox ratio (FLIRR)^48^. Since the fluorescence signatures of NADH and NADPH overlap, they are often referred to as NAD(P)H in literature. NAD(P)H (electron donors) and FAD (an electron acceptor) are metabolic coenzymes in live cells, whose autofluorescence intensity ratio reflects the redox states of the cells and the shifts in the metabolic pathways. However, intensity-based metrics (e.g., ORR) often suffer from wavelength- and depth-dependent light scattering and absorption issues when they are used to characterize the metabolic states of tumor tissues^48^. In contrast, lifetime-based metrics (e.g., FLIRR) bypass these issues, revealing bias-free protein-binding activities of NAD(P)H and FAD. As ORR and fluorescence lifetimes of NAD(P)H and FAD provide complementary information, they have been combined into the OMI index that can distinguish drug-resistant cells from drug-responsive cells in tumor organoids^49^.

Here we demonstrate that *flimGANE* provides rapid and accurate autofluorescence FLIM images of live HeLa cells. DFD data at two emission channels (NAD(P)H: 425-465 nm and FAD:511-551 nm) were collected by the ISS confocal scanning system (with 405 nm excitation) and the acquired data were analyzed by the four methods, generating both intensity and FLIM images (**Fig. 5a-b**). We adopted FLIRR (α_2_NAD(p)H_/α_1_FAD_) as a metric to assess the metabolic response of cancer cells to an intervention. It was found that *flimGANE*-derived FLIRR was highly correlated with its counterpart derived by *TD_MLE* (**Fig. 5c-d; Supplementary Table 13**). Since the NAD(P)H signals came from both the mitochondrial oxidative phosphorylation and cytosolic glycolysis and the FAD signals mainly originated from the mitochondria, image segmentation was often performed to deduce the relative contributions of oxidative phosphorylation and glycolysis to the cellular redox states and help quantify the heterogeneity of cell responses^48^. In our analysis, an intensity threshold was selected to isolate the mitochondrial regions from the rest of the cell area, where the nuclei were manually zeroed (see Methods, **Fig. 5e**). Again, *flimGANE* outperformed the other three methods, generating results most similar to those found in literature^48, 50, 51, 52^, where the peak of FLIRR of cancer cells is usually located at 0.2-0.4 (**Fig. 5f**). *TD_LSE* and *DFD_LSE* provided an incorrect representation, where the former was largely skewed by the low FLIRR values and the latter showed two unrealistic peaks. *TD_MLE* gave a distribution similar to that of *flimGANE*, but with a larger FLIRR peak value, due to the inaccurate estimate of NAD(P)H lifetime under photon-starved conditions.

**Fig. 5.**
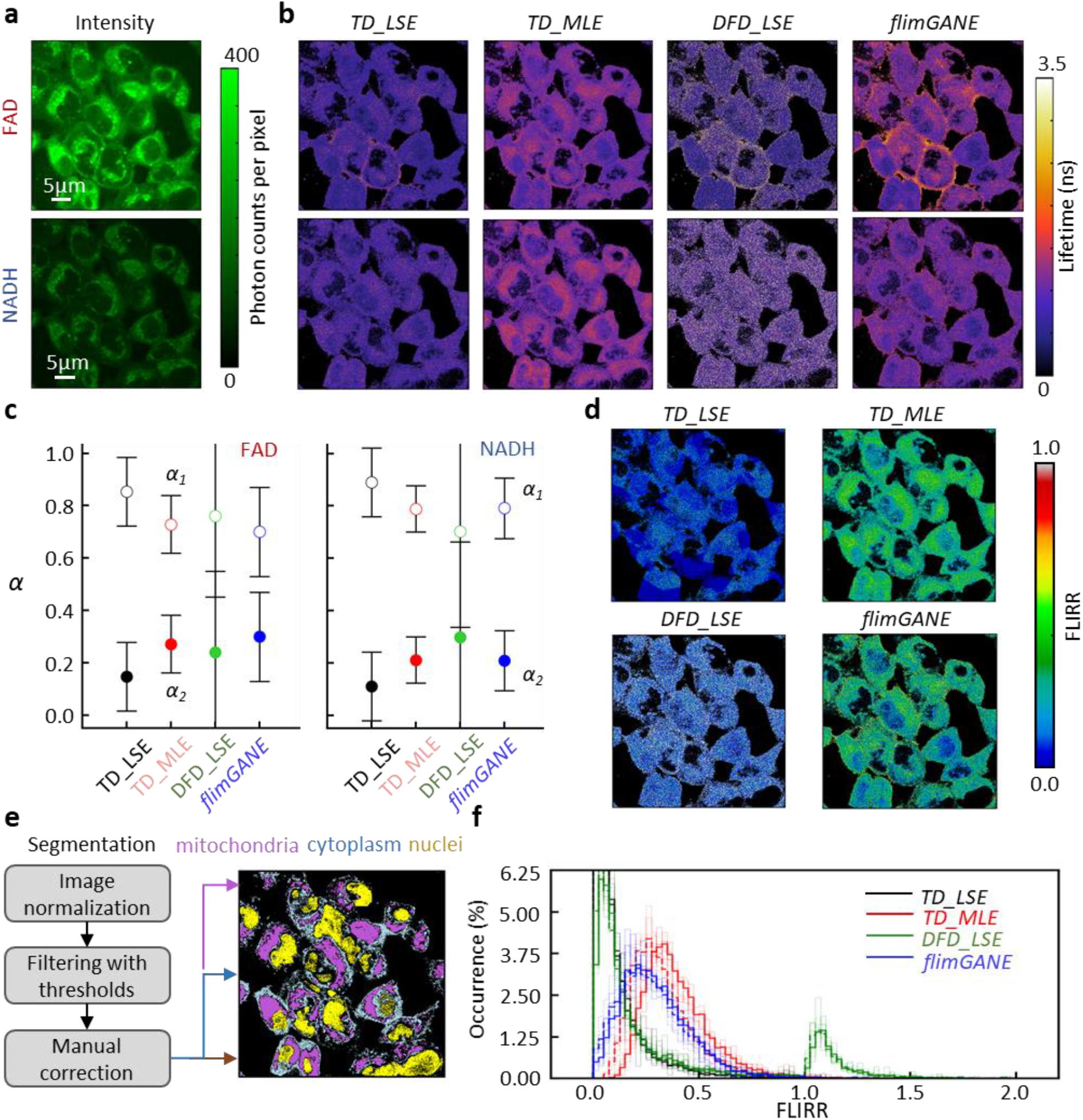
Deep-learning enabled metabolism quantification from low-photon-count autofluorescence FLIM imaging with live HeLa cells. (**a**) Intensity contrast images of FAD and NAD(P)H, respectively. (**b**) FLIM images generated by *TD_LSE, TD_MLE, DFD_LSE* and *flimGANE* of FAD and NAD(P)H, respectively. (**c**) Summary of pre-exponential factor of FAD and NAD(P)H obtained by different methods (error bars, standard deviation, n = 160k pixels). (**d**) FLIRR images showing that *flimGANE*-based quantification matches well with *TD_MLE*. (**e**) Intensity contrasts from (**a**) were normalized for the segmentation of mitochondria, cytoplasm, and nuclei. (**f**) Comparison of FLIRR for redox states obtained from *TD_LSE, TD_MLE, DFD_LSE*, and *flimGANE* (solid line: mitochondria only; dashed line: whole cell analysis except for the nucleus; n = 5 cells).

## Discussion

*flimGANE* addresses an unmet need for FLIM analysis – a computationally efficient, high-throughput and high-quality method for fluorescence lifetime estimation that works reliably even in ultra-low-photon-count conditions (e.g., 50 photon counts per pixel; **Fig. 1d**). In the cases studied above, *flimGANE* generated FLIM images with quality similar to those produced by the gold standard *TD_MLE*, but *flimGANE* clearly outperforms *TD_MLE* in barcode identification (**Fig. 2**), FRET characterization (**Fig. 4**), and metabolic state analysis (**Fig. 5**). We emphasize that in this report we intentionally acquired fluorescence data under low-to medium-light conditions in order to demonstrate the capabilities of the four methods, and we found even the gold standard *TD_MLE* may not necessarily give consistent lifetime estimates under different excitation powers (**Fig. 3g**). It is thus critically important for users to understand the limitations of their lifetime analysis methods, especially when handling the low-count decays. Here we provide an alternative FLIM analysis approach for users to consider, where the low-laser power requirement will reduce photobleaching and phototoxicity issues in delicate samples.

As FLIM finds more clinical applications such as retina imaging^53^ and tumor margin identification^54^ in recent years, it becomes critically important that we have a fast, fit-free and accurate method to perform the lifetime imaging analysis. Whereas the previous deep-learning methods also provided high-speed FLIM analysis^55, 56^ and addressed the low-photon count issues^56, 57^, by using GAN in our model we may gain additional advantage in the image quality over the standard CNN^28^. Our *flimGANE* takes raw fluorescence decay histograms and experimental IRF as inputs and adopts a virtual resampling procedure that is integrated into the model. Through the use of convolutional block that mitigates the artifacts dependent on neighboring temporal bins and residual block that allows for the flow of memory from the input layer to the output layer, our generative model generates high-quality decays based on low-photon-count inputs without introducing bias. The inference of lifetime is non-iterative and does not require parameter search to perfect the network performance. In this work, we evaluated the network performance using *in-silico* data, demonstrating that *flimGANE* can generate reasonable lifetime estimates with photon counts as low as 50 per pixel. However, considering background noises and other imperfect conditions in the real experiments, 100 photons per pixel are likely still required to get a reliable lifetime estimate.

To the best of our knowledge, this is the first demonstration of a GAN model applied to reconstruct FLIM images. Since the use of Jensen-Shannon divergence as the objective function can cause problems such as vanishing gradients and mode collapse during GAN training, we incorporated Wasserstein metric in our model which provides much smoother value space to avoid those issues^31^. We are continuing to explore the incorporation of other frameworks in our model, including the gradient penalty (WGAN-GP)^58^, the sequence generation framework (SeqGAN)^59^, and the context-aware learning^60^, that may in some instances provide more suitable approximate inference.

While *flimGANE* provides rapid, accurate and fit-free FLIM analysis, its cost lies in the network training. In other words, *flimGANE* is particularly valuable for the FLIM applications where retraining is not frequently required. For instance, samples have similar fluorophore compositions (i.e., autofluorescence from metabolites in patient-derived organoids) and IRF of the imaging system is stable and seldom changes. Different training datasets were employed to train the model separately that eventually led to the results shown in **Figs. 1**-**5** (**Supplementary Table 1**). A primary reason to retrain the model is due to the change of IRF (**Supplementary Fig. 14**). Whenever a different laser source is chosen for excitation, the filters are replaced, or the optics system is realigned, the IRF can also change and the network should be retrained. For an entirely new imaging system, it can take more than 500 hours to fully train the network with a lifetime range of 0.1-10.0 ns (two components, *τ*_*1*_ and *τ*_*2*_) and a pre-exponential factor range of 0.0-1.0 (for *α*_*1*_). However, if we know the range of lifetime of interest on our samples (e.g., 1.3-4.0 ns as the two lifetime components for barcode identification and 0.5-5.0 ns for live HeLa cell studies), a smaller training dataset can be employed to speed up the training process (e.g., 19 hours in **Supplementary Table 1**).

Transfer learning^61^ from a previously trained network for another type of sample can also speed up the convergence of the learning process. However, this is neither a replacement nor a required step for the entire training process. After running a sufficiently large number of training iterations for generator (> 2,000), the optimal network can be selected when the validation loss no longer decreases. No matter what kinds of data sets is used for model training, *flimGANE* can rapidly generate batches of FLIM images without using a graphics processing unit (GPU). Notably, an essential step in generating a FLIM image by our network is the accurate alignment of a fluorescence decay with respect to its corresponding IRF. A multi-stage preprocessing step, termed CoME (**Supplementary Fig. 10**), was employed to bypass the instability issues in the TCSPC electronics, leading to bias-free estimates of fluorescence lifetimes.

Taken together, our work represents an important step forward towards real-time and super-resolution^62, 63^ FLIM. In fundamental biological research, further development of *flimGANE* will enable monitoring of fast binding kinetics and molecular dynamics inside live cells. In medicine, *flimGANE* can provide rapid identification of tumor-free margin during tumor surgery^54^ and investigation of disease progression in the retina^53^. We envision that our method will soon replace *TD_MLE* and *TD_LSE* analysis packages in some commercial FLIM systems.

## Materials and methods

### Structure of dataset

The dataset is composed of training data and testing data. Both training and testing data can be obtained either by a Monte Carlo (MC) simulation with the parameters or by acquiring experimental data from the ISS Alba v5 confocal microscopic system. The experimental data is the fluorescence decay histogram matrix of dimension *n* by *n* by *b* (*n* is the image size, which is either 256 or 512; *b* is the number of histogram bin size, which is 256 in this work).

### Structure of *flimGANE*

The *flimGANE* consists of a generator, a discriminator, and an estimator. The generator is composed of a convolutional block, a multi-task layer associated with the rectified linear unit (ReLU) activation function, a decoding layer associated with the tanh activation function, and a residual block implicitly. The discriminator is composed of four layers of neural networks. All the layers are fully-connected layers composed of 128, 64, 8, and 1 node. The first three layers are associated with the sigmoid activation function, and the last one used linear function. The estimator begins with two fully-connected neural networks with 64 nodes for incoming inputs, followed by the concatenation layer and a multi-task layer associated with ReLU activation function (for further details, see Supplementary Methods).

We trained *flimGANE* with three-stage processes: generative model training, estimative model training, and *flimGANE* combination training. In generative model training, we adopt Wasserstein GAN algorithm, where the generator and the discriminator are trained with Wasserstein loss. In estimative model training, the estimator is trained with the mean squared error cost function. In *flimGANE* combination training, well-trained generator and estimator are combined and trained with the mean squared error cost function. The RMSprop optimizer is applied to both the generator and the discriminator by setting the learning rate as 5 × 10^−5^. The Adam optimizer is applied to the estimator by setting the learning rate as 0.001 (for further details, see Supplementary Methods).

### Evaluation metrics

To compare the proposed methods with other existing algorithms, we utilized several metrics, including execution time, mean squared error for pixel-wise comparison, peak signal-to-noise ratio, structural similarity index, and visual information fidelity for the quality of the FLIM image with respect to the reference FLIM image (for further details, see Supplementary Methods).

## Acknowledgements

The authors thank Mr. Li-Heng Chen from Prof. Alan C. Bovik’s Image & Video Engineering (LIVE) lab at University of Texas at Austin and Dr. Beniamino Barbieri from ISS for discussion and suggestions. This work is supported by NSF (CHE1611451 to H.-C.Y.), the Welch Foundation (Grant F-1833 to H.-C.Y.), NIH (Grant GM129617 to H.-C.Y.), Texas 4000 foundation and CPRIT grant RR160005 (to T.E.Y.). T.E.Y. is a CPRIT Scholar in Cancer Research. Y.-I.C. is supported by the University Graduate Continuing Fellowship at UT Austin.

## Author contributions

Y.-I.C., Y.-J.C. and H.-C.Y. conceived the project and wrote the article. Y.-I.C., Y.-J.C. and S.-C.L. developed the image processing and analysis software. Y.-I.C., J.Y. and H.-C.Y. designed the experiments. Y.-I.C., J.Y., Y.-A.K., and S.H. prepared samples, performed cell culture, and collected images. S.-C.L., T.D.N. supported the special instrumental setup for all experiments. H.G.R., S.R.S., T.E.Y. and Y.-L.L. advised the experimental design and data analysis. H.-C.Y. supervised the project.

