## Supplementary file for "Deep learning enables rapid and robust analysis of fluorescence lifetime imaging in photon-starved conditions"

**Supplementary Methods**

**Supplementary Figures**

Supplementary Fig. 1 Generation of the simulated decay histogram with lifetime parameters and photon counts by MC simulation

Supplementary Fig. 2 flimGANE, a fluorescence lifetime analysis algorithm, can generate accurate FLIM images with well-trained three subnets, generator (G), discriminator (D), and estimator (E)

Supplementary Fig. 3 Generator (G), a subnet from flimGANE, can transform low-photon-count decay histogram into high-photon-count one

Supplementary Fig. 4 flimGANE outperformed other analysis methods in FLIM images reconstruction at various photon-count conditions in-silico

Supplementary Fig. 5 The performance of different analysis methods was evaluated by MSE and SSIM

Supplementary Fig. 6 The performance of different analysis methods was evaluated by quality index (QI)

Supplementary Fig. 7 Difference of center of mass (CoM) between the IRF and decay histogram was demonstrated positively correlated

Supplementary Fig. 8 Calibration line for center of mass evaluation (CoME) was consistent over different photon counts

Supplementary Fig. 9 A real-world application of the CoME

Supplementary Fig. 10 flimGANE can accurately discriminate fluorescence lifetime barcode beads individually at low-photon-count condition

Supplementary Fig. 11 Two dimensional (lifetime versus intensity) scattered plots from different methods illustrated the success of flimGANE for identifying three populations of barcodes with nearly the same size
Supplementary Fig. 12 CFP-g-YFP-transfected cells were selected from fluorescence image by ImageJ (ROI Manager)
Supplementary Fig. 13 The quantification of CFP-g-YFP-transfected MDA-MB-231 cell FLIM images for YFP channel
Supplementary Fig. 14 Incorrect Convallaria FLIM image was reconstructed by flimGANE with incorrect IRF input

**Supplementary Tables**

Supplementary Table 1 Training database for different applications
Supplementary Table 2 Mean squared error (MSE) comparison of different analysis methods in silico
Supplementary Table 3 Peak signal-to-noise ratio (PSNR) comparison of different analysis methods in silico
Supplementary Table 4 Structural similarity index (SSIM) comparison of different analysis methods in silico
Supplementary Table 5 Goodness-of-fit ($\chi 2$) comparison of different analysis methods in silico
Supplementary Table 6 Execution time comparison of different analysis methods in silico
Supplementary Table 7 Summary of different analysis methods for the mixture of two fluorescent dyes
Supplementary Table 8 Summary of each detected bead
Supplementary Table 9 MSE, PSNR, SSIM, and VIF comparison of different analysis methods for Convallaria FLIM images (Standard: medium-count TD_MLE FLIM)
Supplementary Table 10 MSE, PSNR, SSIM, and VIF comparison of different analysis methods for live HeLa cell FLIM images (Standard: medium-high-count FLIM)
Supplementary Table 11 MSE, PSNR, and SSIM comparison of different analysis methods for FRET FLIM images (Reference: TD_MLE FLIM)
Supplementary Table 12 Apparent lifetime of CFP calculated with Gaussian distribution fitting
Supplementary Table 13 MSE, PSNR, and SSIM comparison of different analysis methods for autofluorescence FLIM images (Reference: TD_MLE FLIM)

### Supplementary Methods

**Generative adversarial network structure and training.** In this work, we trained a deep neural network using a generative adversarial network (GAN)[1]. GAN consisted of two subnets, generator (*G)* and a discriminator (*D)*, which were trained via an adversarial process. *G* mimicked the ground-truth fluorescence decay histogram while *D* returns an adversarial loss to the *G*-output fluorescence decay, as illustrated in **Supplementary Fig. 2** The cost functions for the gradient descent update of generator ($G_{cost}$) and discriminator ($D_{cost}$) were designed as follows, respectively.

$$G_{cost} = \frac{1}{n}\sum_{i=1}^{n} log\left(1 - D\big(G(z_i)\big)\right) \tag{1}$$

$$D_{cost} = \frac{1}{n}\sum_{i=1}^{n}\left(\log\big(D(x_i)\big) + \log\left(1 - D\big(G(z_i)\big)\right)\right) \tag{2}$$

where $z_i$ represents the normalized low-photon-count fluorescence decay histogram, and $x_i$ is the normalized ground-truth fluorescence decay histogram. $G(z)$ is the normalized ground-truth mimicking histogram ($G_{output}$), and $D(x)$ represents the probability that $x$ came from the ground-truth fluorescence decay histogram rather than $G_{output}$.

While GAN had shown great success, there exist several problems, including the difficulty to achieve Nash equilibrium[2] vanish gradient due to the use of loss functions in equation[3] (1, 2). Wasserstein generative adversarial network (WGAN)[4] was used to improve the *G* performance. Wasserstein distance instead of Jensen-Shannon divergence was used to train GAN by evaluating the distance of the distributions between the real data and the generated data. The improved training process could offer strong enough gradients to train the generator than that of the original GAN. The cost functions of WGAN for the gradient descent update of *G* and *D* were as follows,

$$G_{cost} = -\frac{1}{n}\sum_{i=1}^{n} f\big(G(z_i)\big) \tag{3}$$

$$D_{cost} = \frac{1}{n}\sum_{i=1}^{n}\Big(f(x_i) - f\big(G(z_i)\big)\Big) \tag{4}$$

$$|f(x_1) - f(x_2)| \le |x_1 - x_2| \tag{5}$$

Where $f(x)$ is a 1-Lipshitz function which is defined in equation (5). Higher value of *D* output refers to the ground-truth data; while lower value refers to the low-photon-count histogram. This modification of loss function helps to stabilize the training schedule and ensure that training process can lead the deep learning model to converging.

**Generative model (*G*).** Residual neural network[5] and convolution neural network are artificial neural networks (ANNs) architecture that were first proposed for image recognition, easing the training of networks that are substantially deeper than those used previously. A similar network architecture has also been successfully applied in recent image reconstruction and image synthesis applications[6]. The structure of the generative network used in this work is demonstrated in **Fig. 1** and **Supplementary Fig. 2**, which consists of a convolutional block, a multi-task layer (n = 3), a decoding layer explicitly, and a residual block implicitly. Inside the residual block, the convolutional block consists of two convolutional layers and pooling layers followed by a flatten layer, within which it performs

$$y = \mathrm{ReLU}[\mathrm{Conv}\{\mathrm{ReLU}[\mathrm{Conv}\{\mathrm{Concat}(x_{1:256}, x_{256:512})\}]\}] \quad (6)$$

where $x$ represents the input of the generative model, normalized low-photon-count fluorescence decay histogram and corresponding instrument response function (IRF), $y$ is the output of the convolutional block. $\mathrm{Concat}()$ is the concatenation operation of two inputs. $\mathrm{Conv}\{\}$ is the convolution operation, $\mathrm{ReLU}[]$ is the rectified linear unit activation function,

$$\mathrm{ReLU}[x] = x^{+} = \max(0, x) \quad (7)$$

The dimension of the output of each ReLU activation function is reduced by AveragePooling layer. Then a multi-task neural network with hard parameter sharing converts the high dimensional flattened output into three tasks, and each task corresponds to each lifetime parameter (for example, bi-exponential decay model). The last network, decoding layer, termed multilayer perceptron with the activation functions as tanh() to force the range of the output to lie within -1 and 1, maps the 3 tasks into 256 channels of output that corresponds to the fluorescence decay histogram. Instead of learning a direct mapping toward ground-truth fluorescence decay histogram, we reframe the process with the residual learning framework by introducing a residual connection between one of the inputs, normalized low-photon-count decay histogram, and the model's output.

**Discriminative model (*D*).** As shown in **Supplementary Fig. 2**, the structure of the discriminative model begins with a densely connected neural network with 128 nodes. Then the output is fed into other densely-connected neural networks with 64, 8, and 1 node. All layers instead of the last one has a sigmoid activation function, which forces the output to lie within the range from 0 to 1, defined as follows,

$$\mathrm{sigmoid}(x) = \frac{1}{1+e^{-x}} \quad (8)$$

The last layer has the linear activation function to output the score corresponding to the input histogram fed into *D*.

**Estimator model (*E*).** On top of the components in the GAN framework, here we also constructed an estimator that focused on predicting the lifetime parameters of interest from the high-photon-count fluorescence decay histogram. In *flimGANE* analysis, the output of the generator, normalized ground-truth mimicking fluorescence decay histogram, would be fed into the estimator model, *E*, to infer the lifetime values. This hold a great potential to improving the predictive power of the lifetime for more than two species. As shown in **Supplementary Fig. 2**, the structure of the estimator model begins with two densely connected neural networks with 64 nodes for incoming IRF and high-photon-count fluorescence decay histogram, respectively. These two outputs are concatenated by a concatenation layer. The output of the concatenation layer is fed into the multi-task neural network (n = 3) with hard parameter sharing, and a multilayer perceptron with a single hidden layer, whose output is the corresponding fluorescence lifetime parameters. The loss function for *E* to be trained is defined as follows,

$$E_{loss} = \frac{1}{n}\sum_{i=1}^{n}(y_i - \hat{y}_i)^2 \quad (9)$$

Where $y_i, \hat{y}_i$ represent the predicted and the ground-truth lifetime parameters, respectively.

**Network training schedule.** During our training the batch size is set as 32 on single GPU. In this work, we employed three-stage training: generative model training stage, estimative model training stage, and *flimGANE* combination training stage. In generative model training stage, total iteration was set to be 2,000. Within each iteration, we randomly selected ~3% training samples from the pool of dataset. The discriminative model was updated five times while the generative model was kept untrainable, then the generative model is updated once while keeping the other one untrainable. In estimative model training stage, total iteration was set to be 500. Within each iteration, we randomly selected ~18% samples (90% for training, 10% for validation) from the pool of dataset. The estimator model was then updated ten times. The aforementioned two training stages can be trained simultaneously and independently. Last, in *flimGANE* combination training stage, we combined the generative model and estimative model together, and fed *flimGANE* with randomly selected 10% training samples to update estimative model only for 100 times in each iteration, where the total iteration was set to be 100. Both the generative model and discriminative model were randomly initialized by Glorot uniform initializer and optimized using RMSprop optimizer with a starting learning rate of $5 \times 10^{-5}$. The final generative and discriminative models for each application in this work were selected at around 2000$^{th}$ iteration, which took 1.5h to train in generative model training stage. The estimator model which maps ground-truth fluorescence decay histogram to lifetime values was initialized by Glorot uniform initializer and optimized using Adam optimizer[7] as well with learning rate of $1 \times 10^{-3}$. All the estimative models for specific applications were pre-trained for 500 iterations, which took ~8 minutes to train in estimative model training stage. The generative models integrated with estimative models were then trained for 100 iterations, which took ~35 minutes to train in *flimGANE* combination training stage. Training without the discriminative loss and predictive cost can result in over-smoothed images, as the generative model optimizes only a specific group of statistical metrics. Therefore, it is imperative to incorporate discriminator to train the generative model well. A step-by-step training instruction and guideline, with several critical steps discussed and emphasized, are provided in **Supplementary Fig. 2**.

**Software and hardware used for *flimGANE* development.** This framework was implemented with TensorFlow framework version 1.12.0 and Python version 3.6.5 in the Microsoft Windows 10 operating system. The training was performed on a custom-built desktop equipped with single GeForce RTX 2070 XC GPU cards (NVIDIA) and a Core i7-9700K CPU @ 3.6 GHz (Intel).

**Generation of the simulated database (*in silico*) with Monte Carlo (MC) method.** First, multiple sets of ground truth are determined based on the lifetimes ($\tau_1$ and $\tau_2$) and the fraction amplitude ($\alpha_1$). For each ground truth, different photon counts (*pcs*) and number of duplicates (e.g., 100) are assigned to construct the training dataset. For every training sample, it was assigned a value of short lifetime ($\tau_1$), long lifetime ($\tau_2$), fraction amplitude of short lifetime species ($\alpha_1$), and photon counts (*pcs*). The IRF is obtained by averaging across all the pixels of the calibration image taken at the beginning of the experiment. These parameters were employed to generate the probability mass function that describes the distribution of the photon arrival time via the equation (10):

$$P(t) = N\left(IRF(t) \otimes \left[\alpha_1 e^{-\frac{t}{\tau_1}} + (1-\alpha_1) e^{-\frac{t}{\tau_2}}\right]\right)$$
$$N\big(f(t)\big) = \frac{f(t)}{sum\big(f(t)\big)} \quad (10)$$

Given the probability mass function, we can perform Monte Carlo simulation method to extract specified number (photon counts, *pcs*) of samples. Those extracted samples were then used to generate the simulated (degraded) decay histogram.

**Fluorescence lifetime imaging microscopy (FLIM).** The FLIM system were equipped with diode lasers providing 370 nm, 405 nm, 488 nm, and 635 nm lines and a supercontinuum white laser (SuperK EVO, NKT Photonics) providing a wavelength within 400 – 1000 nm. All the laser lines combined in Alba v5 passed a multi-band dichroic mirror to excite samples. Fluorescence emission light came back along the same path to the multi-band dichroic mirror, and went into 3 detectors, where each detector has its own pinhole. A Nikon inverted microscope with 60x, NA 1.2 water objective is used. An ASI XY automatic stage with motorized Z control is equipped the current setup. The photon counts were acquired with fastFLIM unit to build up the phase histogram. Here the laser repetition period we used is 50 ns, and it is divided into 256 bins.

**Correction of the temporal shift between instrument response function (IRF) and acquired fluorescence decay histogram.** Temporal shift occurred when the measurement is biased toward shorter or longer arrival time due to the noise. To characterize the temporal shift, the most common method is to perform model fitting with time shift as the additional parameter. During the process of curve fitting, the algorithms take temporal shift into consideration to find the optimal value for both lifetime parameters and time shift. However, this could take a large amount of time to obtain the optimal time shift parameter. Here we introduced a new analysis, Center of Mass Evaluation (CoME), to quantify the time shift parameter without the time-consuming curve fitting process. The mass center of the histogram, also known as the expected value of the histogram, was determined by the following equation,

$$CoM = \frac{\sum x_i h(x_i)}{\sum h(x_i)} \quad (11)$$

where $h(x_i)$ is the fluorescence decay histogram acquired from the experiment at the corresponding bin $x_i$.
We hypothesized that for certain lifetime, the difference of CoM between IRF and the fluorescence decay histogram should be the same. To verify the above hypothesis, we generated 250 simulated decay histograms under 150, 500, 1500, 5000 photon counts for various fluorescence lifetime values and calculated the mean and the standard deviation of the difference of CoM. **Supplementary Fig. 13** shows the calibration line for the distance of CoM between IRF and fluorescence decay histogram. Such a calibration line is employed to determine the amount of the temporal shift.

**Implementation of center of mass evaluation (CoME) and trained *flimGANE*.** To eliminate the "temporal shift" effect that may cause inaccurate lifetime estimation, this parameter was obtained by "center of mass" analysis with the control experiment and then employed to calibrate the measurement for each pixel. With IRF and the calibrated fluorescence decay histogram as model inputs, *flimGANE* outputs two lifetime components and its relative ratio for each pixel, forming the FLIM image accurately and rapidly.

**Fluorescence lifetime fitting with time-domain least-squares estimation (*TD_LSE*).** Assume the fluorescence decay histogram contains either only one or two species:

$$f_{one\ species}(t, \alpha_1 = 1, \tau_1, \tau_2 = 0) = f(0)\ e^{-t/\tau_1} \tag{12}$$

$$f_{two\ species}(t, \alpha_1, \tau_1, \tau_2) = f(0)\ \left(\alpha_1 e^{-\frac{t}{\tau_1}} + (1-\alpha_1)e^{-\frac{t}{\tau_2}}\right) \tag{13}$$

The expected measurement is the convolution of the fluorescence decay histogram and the IRF,

$$m(t, \alpha_1, \tau_1, \tau_2) = IRF(t) \otimes f(t, \alpha_1, \tau_1, \tau_2) \tag{14}$$

where $\tau_1$ represents short lifetime, $\tau_2$ represents long lifetime, $\alpha_1$ is fraction amplitude of short lifetime species, and *IRF* is obtained from the experiments. The best fit parameters are determined by minimizing the sum of squared error

$$R^2 = \sum[y_i - m(t_i, A, \tau_1, \tau_2)]^2 \tag{15}$$

where *y* is the data to be fitted.

**Fluorescence lifetime fitting with time-domain maximum likelihood estimation (*TD_MLE*).** The observed fluorescence decay histogram is given by equation (14), where $IRF$ represents the instrument response function, $\tau_1, \tau_2$ are the shorter and longer fluorescence lifetime, and $\alpha_1$ is the fraction amplitude of shorter lifetime. The goal of MLE is to find the parameters that maximize the likelihood function, which is equivalent to minimizing the negative likelihood function. For the case of fluorescence lifetime analysis, we aim to minimize the negative Poisson log-likelihood function (-*L*), which is defined as follows,

$$-L = \sum_{i=0}^{N-1}(m_i - n_i log m_i + log n_i!) \tag{16}$$

where, $n_i$ represents the number of the detected photons in $i^{th}$ bin and $m_i$ represents the expected measurement in $i^{th}$ bin.

**Fluorescence lifetime fitting with digital frequency-domain least-squares estimation (*DFD_LSE*).** Assuming an infinite short excitation pulse ($\delta$-function), fluorescence decay histogram composed of *N* ($N \geq 1$) fluorescent species with distinct fluoresce lifetimes can be modeled and written in a general form as follows,

$$f(t) = f(0) \sum_{i=1}^{N} \alpha_i e^{-t/\tau_i} \tag{17}$$

Where, *f(0)* is the number of the instantly emitted photons at time 0; the coefficient $\alpha_i$, called the pre-exponential factor, is the fraction amplitude and $\tau_i$ is the fluorescence decay time of the *i*-th component of the mixture. The amplitude weighted lifetime $\tau_\alpha$ (often called apparent lifetime) is given by,

$$\tau_\alpha = \sum_{i=1}^{N} \alpha_i \tau_i \tag{18}$$

The average lifetime $\tau_f$ is given by the equation (19), where the fraction $f_i$ is weighted in consideration of the $i^{th}$ lifetime ($\alpha_i \tau_i$).

$$\tau_f = \sum_{i=1}^{N} f_i \tau_i$$
$$f_i = \frac{\alpha_i \tau_i}{\sum_{i=1}^{N} \alpha_i \tau_i} \tag{19}$$

Due to the finite response of a system, the measured decay signal *f'(t)* is the convolution of the fluorescence decay histogram and the IRF plus the noise, *n(t)*,

$$f'(t) = IRF \otimes f(t) + n(t) = IRF \otimes \left\{ f(0) \sum_{i=1}^{N} \alpha_i e^{-t/\tau_i} \right\} + n(t) \tag{20}$$

Thus, accurate FLIM data analysis typically requires the calibration of the IRF. In time-domain FLIM, the IRF can be measured by recording the scattered excitation light when using one-photon excitation, or the second-harmonic generation (SHG) signals for two-photon excitation by using a sample that yields strong SHG, such as urea crystal. In DFD-FLIM, the IRF is typically calibrated with a known fluorescence lifetime standard.

In TCSPC FLIM, a decay histogram *f'(t)* is recorded at each pixel location. To present the data in frequency domain, we take the Fourier Transform of the fluorescence decay histogram to get real and imaginary components.

$$\vec{F}(\omega) = \mathcal{F}\big(f'(t)\big) = \mathcal{F}\big(IRF \otimes f(t) + n(t)\big) = \mathcal{F}(IRF) * \mathcal{F}\big(f(t)\big) + \mathcal{F}\big(n(t)\big) \tag{21}$$

Notice that in frequency domain, the convolution with the IRF is a simple multiplication of the IRF vector, and the noise is a simple addition of the noise vector. During the calibration procedure, we simply subtract the noise vector and divide the denoised vector by the IRF vector to get the emission fluorescent vector. If the emission fluorescent vector is from the standard calibration single component dye, the phasor will be on the semi-circle after the calibration. The later measurement of the unknown samples will be always noisy and IRF-free for finding the unknown fluorescent lifetime(s). The Fourier Transform of the fluorescence decay histogram can be separated into real number, *g(ω)*, and imaginary number, *s(ω)*.

$$g(\omega) = \frac{\int_0^\infty f'(t) cos(\omega t)}{\int_0^\infty f'(t)} \tag{22}$$

$$s(\omega) = \frac{\int_0^\infty f'(t) sin(\omega t)}{\int_0^\infty f'(t)} \tag{23}$$

Therefore, after calibration, each fluorescent decay histogram (without IRF and noise) of each pixel can be plotted as a single point, termed phasor, in the phasor plot by applying the sine and cosine transforms to the measured decay data, where the modulation frequency $\omega$ is the laser repetition angular frequency and is calculated by multiplying the laser repetition rate with 2π.

By solving the integral, we can then derive the following relationships between the phasor and the lifetime,

$$g(\omega) = \sum_{i=1}^{N} \frac{f_i}{1+\omega^2 \tau_i^2} \tag{24}$$

$$s(\omega) = \sum_{i=1}^{N} \frac{f_i \omega \tau_i}{1+\omega^2 \tau_i^2} \tag{25}$$

where $N$ is the number of the fluorescent species, and $f_i$ is the fractional contribution of the $i^{th}$ species with the fluorescence lifetime $\tau_i$.

**Reduced chi-square ($\chi^2$) for time-domain and digital frequency-domain data.** In time-domain fitting process, the goodness-of-fit is evaluated by the reduced $\chi^2$, which is defined as

$$\chi^2 = \frac{1}{n-p-1}\sum_{k=1}^{n}\frac{[N(t_k)-N_c(t_k)]^2}{N(t_k)} \tag{26}$$

where $n$ is the number of acquisition bin, $p$ is the number of the fitting parameters, $N(t_k)$ is the number of photons collected at bin $k$, and $N_c(t_k)$ is the number of photons at bin $k$ calculated from the fitting parameters and the decay model.

The reduced $\chi^2$ for digital frequency-domain fitting is defined as

$$\chi^2 = \frac{1}{2n-p-1}\left\{\sum_{k=1}^{n}\left[\frac{\varphi_k-\varphi_{ck}}{\sigma_{\varphi k}}\right]^2 + \sum_{k=1}^{n}\left[\frac{m_k-m_{ck}}{\sigma_{mk}}\right]^2\right\} \tag{27}$$

where $n$ is the number of modulation frequencies, $p$ is the number of the fitting parameters, $\varphi_k$ and $m_k$ are the measured phase shift and modulation ratio at frequency $k$, $\varphi_{ck}$ and $m_{ck}$ are the calculated phase shift and modulation ratio at frequency $k$ using the fitting parameters and the selected model, $\sigma_{\varphi k}$ and $\sigma_{mk}$ are the uncertainties in the phase shift and modulation ratio values at frequency $k$, respectively.

**Pre-processing of FLIM data.** Calibrating the phasor plot is the only procedure for digital frequency domain fitting. The DFD-FLIM data measurements at each pixel location are composed of both the phase delay ($\varphi$) and the amplitude modulation ratio ($m$). The DFD-FLIM data at each pixel can be mapped to a single point called "phasor" in the phasor plot through a transform defined below, where $\omega$ is the modulation frequency, $g(\omega)$ and $s(\omega)$ represent the values at the two coordinates ($g(\omega), s(\omega)$) of the phasor plot.

$$g(\omega) = mcos(\varphi) \tag{28}$$

$$s(\omega) = msin(\varphi) \tag{29}$$

In order to establish the correct scale for the phasor analysis, the coordinates of the phasor plot need to be calibrated using a standard sample of known lifetime. This will include the calibration of the IRF and the background noise. In DFD-FLIM, this is done during experimental calibration prior to the data acquisition. There is no need to measure the IRF explicitly. The calibration procedure subtracts the noise and divides denoised data by the IRF to reveal the true fluorescent emission component(s). In time domain FLIM, a directly recorded IRF data for the zero lifetime (scatter) will be measured. The IRF will then be used for lifetime fitting, with a convolution involved, and model analysis.

**Post-processing of FLIM (intensity) image.** Either a median or Gaussian filter can be applied before analyzing the data in phasor plot. The degree can be chosen from 0 to 10. When degree is 0, no filter is used. When the degree is 1, it runs through all the pixels once with a 3x3 filter. When degree is 2, it runs through all the pixels twice with a 3x3 filter.

For lifetime fitting, we can bin the data of each pixel to reduce the uncertainty. If the bin number is 0, it is the single pixel histogram or phase and modulation. If the bin number is 1, it borrows left, right, up and down 1 pixel, so total is 3x3 pixels as the selected pixel is at the center. In this work, the bin number is set to be 0.

**Calibration of a FLIM system.** Before imaging biological samples, it is necessary to calibrate the FLIM system using a fluorescence lifetime standard. The fluorescence lifetime for many fluorophores has been established under standard conditions, and any of these probes can be used for the calibration of the FLIM system. Since the fluorescence lifetime of a fluorophore is sensitive to its environment, it is critical to prepare the standards according to the conditions specified in the literature, including the solvent and the pH. It is also important to choose a standard fluorophore with excitation, emission, and fluorescence lifetime properties that are similar to those of the fluorophore used in the biological samples. For example, the dye, Coumarin 6, dissolved in ethanol (peak excitation and emission of 460 and 505 nm, respectively), with a reference lifetime of ∼ 2.5 ns, is often used as the calibration standard for the CFPs. It is important to note that if the excitation wavelength is changed, it is necessary to recalibrate with another appropriate lifetime standard.

**Preparation of fluorescence lifetime barcode beads and FLIM setup.** The oligonucleotide-coated microbeads preparation was carried out using a modified protocol recommended by Bangs Laboratories. Briefly, 2 μL (10 mg/mL) streptavidin-coated microbeads (CP01005, Bangs Laboratories Inc) were transferred into a 1.5 mL centrifuge tube. The microbeads were washed with 20 μL 1X PBS twice by centrifuging at 10K rpm for 3 min and resuspended in 1X PBS. The different ratios of mixed biotinylated single-strand DNA probes (**Probe1**: 5'Atto633-TGGTCGTGGGGCAACTGGGTT-biotin (3.5 ns) and **Probe2**: 5'Cy5-TTTTTTTTTTTT-biotin (1.9 ns) were purchased from Integrated DNA Technologies) were added and incubated for 15 min at room temperature with gentle mixing. The coated microbeads were than separated by centrifuging at 10K rpm for 3 min. The unbound biotinylated probes were removed by washing three times in 1X PBS. Then, two species coated beads are now ready for downstream applications. Here we demonstrated three different barcode beads (**Table S6**) imaged by the FLIM. The FLIM images for fluorescence lifetime barcode beads were taken by the laser light focused through a 60X NA=1.2, water immersion objective (CFI Plan Apochromat Lambda 60XC, Nikon). A diode laser was used as an excitation source at 635 nm. The fluorescence was detected with an avalanche photodiode (SPCM-AQR-15, Perkin Elmer) after passing through a bandpass filter (685/40 nm, Semrock). FLIM images (512×512 pixels) were scanned three times with dwell time 0.04 ms/pixel. Cy5 in water (1 ns) was used for calibrating FLIM system.

**Convallaria cover slide and FLIM setup.** The convallaria (lily of the valley) cover slide was stained on 26mm x 76mm glass slides (RE-Cal01, Medical and Science Media). A supercontinuum white laser was used as an excitation source at 630/38 nm. The fluorescence was detected with an avalanche photodiode (SPCM-AQR-15, Perkin Elmer) after passing through a bandpass filter (685/40 nm, Semrock). The FLIM images were taken by the laser light focused through a 60X NA=1.2, water immersion objective (CFI Plan Apochromat Lambda 60XC, Nikon). FLIM images (512×512 pixels) were scanned one time (for low-photon-count condition) and three times (for medium-photon-count condition) with dwell time 0.1 ms/pixel. Cy5 in water (1 ns) was used for calibrating FLIM system.

**Preparation of staining of live cells and FLIM setup.** Live HeLa cells were seeded onto optical imaging 8-well Lab-Tek chambered cover glass (154534, Thermo Fisher Scientific) with cell density 70–90% confluent per well and grown overnight at 37ºC in a humidified atmosphere with 5% $CO_2$ prior to staining. Cells were maintained in

DMEM/F12 medium (11320082, Thermo Fisher Scientific) supplemented with 10% heat-inactivated fetal bovine serum (16140071, Thermo Fisher Scientific) and 50 U/mL penicillin-streptomycin (15070063, Thermo Fisher Scientific). CellMask™ Green (C37608, Thermo Fisher Scientific) or CellMask™ Red (C10046, Thermo Fisher Scientific) plasma membrane stain (1μg/mL) were used to stain the plasma membrane of live cells for 10 min at 37ºC. The staining solution was removed from the chambered cover glass. Then, the live cells were washed with PBS three times. The nucleus was stained by the permeable Hoechst 33342 dye (62249, Thermo Fisher Scientific) for 10 min at 37ºC and washed with PBS three times. Cells were then kept in the phenol red-free DMEM/F12 (21041025, Thermo Fisher Scientific) for the FLIM images acquisition. A diode laser was used as an excitation source at 405, 488 and 635 nm. The FLIM images were taken by the laser light focused through a 60X NA=1.2, water immersion objective (CFI Plan Apochromat Lambda 60XC, Nikon). The fluorescence was detected with an avalanche photodiode (SPCM-AQR-15, Perkin Elmer) after passing through a bandpass filter (494/34, 531/40, 685/40 nm, Semrock). FLIM images (512 × 512 pixels) were taken with dwell time 0.1 and 0.2 ms/pixel, respectively. Alexa 405 in water (3.6 ns), rhodamine 110 in water (4 ns) and Cy5 in water (1 ns) was used for calibrating FLIM system.

**Preparation of intramolecular FRET glucose sensor.** Triple negative breast tumor cell line, MDA-MB-231, was obtained from the American Type Culture Collection (ATCC) and grown in high-glucose (25 mM) DMEM/F12 culture medium (11320082, Thermo Fisher Scientific) containing 10% heat-inactivated fetal bovine serum (16140071, Thermo Fisher Scientific) and 50 U/mL penicillin-streptomycin (15070063, Thermo Fisher Scientific). The plasmid carrying the glucose FRET sensor, pcDNA3.1 FLII12Pglu-700uDelta6 (a gift from Wolf Frommer), was obtained from Addgene. Prior to transfection, MDA-MB-231 cells were seeded in a 6-well plate and allowed with cell density 70–90% confluent per well. Transfections were performed using Lipofectamine™ LTX and Plus™ reagent (15338100, Thermo Fisher Scientific) according to manufacturer's instructions. Transfection medium, Opti-MEM™ I Reduced Serum Medium (11058021, Thermo Fisher Scientific), contained no serum or antibiotics. Six hours post-transfection, the medium was replaced with DMEM culture medium. Three days post-transfection, medium was replaced with DMEM containing 100 μg/mL G418 (A1720, Sigma-Aldrich) for selection. After two weeks of selection, the cells were sorted by flow cytometry (FACSAria, BD Biosciences) based on YFP expression. MDA-MB-231 cells transfected with the FRET glucose sensor were seeded onto optical imaging 8-well Lab-Tek chambered cover glass (154534, Thermo Fisher Scientific) with cell density 70–90% confluent per well and grown overnight at 37ºC in a humidified atmosphere with 5% CO2. The medium was replaced with glucose-free DMEM culture medium for 24 hours before FLIM image acquisition. The FLIM images were taken by the laser light focused through a 20X objective (UPLSAPO, Olympus). The fluorescence of CFP and YFP were detected by two avalanche photodiodes (SPCM-AQR-15, Perkin Elmer) after passing through the bandpass filters (494/34, 585/40 nm, Semrock), respectively. FLIM images (256 × 256 pixels) were scanned three times with dwell time 0.1 ms/pixel. Alexa 405 in water (3.6 ns) was used for calibrating FLIM system.

**Autofluorescence FLIM for cellular metabolism and FLIM setup.** Live HeLa cells were seeded onto optical imaging 8-well Lab-Tek chambered cover glass (154534, Thermo Fisher Scientific) with cell density 70–90% confluent per well and grown overnight at 37°C in a humidified atmosphere with 5% $CO_2$. Before taking an autofluorescence FLIM image, the medium was replaced with phenol red-free complete medium. A diode laser was used as an excitation source at 405 nm. The FLIM images were taken by the laser light focused through a 60X NA=1.2, water immersion objective (CFI Plan Apochromat Lambda 60XC, Nikon). The autofluorescence of NAD(P)H /FAD and were detected by two avalanche photodiodes (SPCM-AQR-15, Perkin Elmer) after passing through the bandpass filters (445/40, 531/40 nm, Semrock), respectively. FLIM images (512×512 pixels) were scanned one time with dwell time 0.1 ms/pixel. Alexa 405 in water (3.6 ns) was used for calibrating FLIM system.

**Intensity contrast segmentation for mitochondria in Autofluorescence FLIM.** Intensity contrasts of both FAD and NAD(P)H images were normalized to the scale, 0 – 1. Normalized value for each pixel are determined by the following equations:

$$\tau_{normalized} = \frac{\tau_{original} - \tau_{min}}{\tau_{max} - \tau_{min}} \tag{30}$$

Given both normalized images, the locations of segmented mitochondrial were verified where the value presenting the specific location was both greater than the threshold. In this work, the threshold was set to be 0.25. The locations of segmented cytoplasm were determined between 0.16 and 0.25. The locations of segmented nuclei were determined between 0.06 and 0.16.

**Quality index (QI) quantification for simulated decay histogram.** To calculate the QI for the fluorescence decay histogram, we assume the amounts of signals is the total photon counts, and the amount of noise is the deviation of fluorescence decay histogram from the ground-truth decay histogram. The relationship can be described as the following equations:

$$QI = \frac{S}{\sum_i |y_i - r_i|} \tag{31}$$

where $S$ represents the total photons of the fluorescence decay histogram, $y_i$ is the fluorescence decay histogram on $i^{th}$ bin, and $r_i$ represents the ground-truth fluorescence decay histogram on $i^{th}$ bin.

**Structural similarity index (SSIM)[8]**. SSIM is a method used for measuring the similarity between two images. First, two images were smoothed with Gaussian filter (σ = 1.5 pixels). Then SSIM index was obtained for each window of a pair of sub-images. It was calculated for squared windows, centered at the same pixel (dx, dy) of two images. The length of a side of the square window was eleven pixels. The SSIM *(x, y)* was then obtained for two windows in two images (*x*, and *y*) as follows,

$$\mathrm{SSIM}(x, y) = \frac{(2\mu_x\mu_y + c_1)(2\sigma_{xy} + c_2)}{(\mu_x^2 + \mu_y^2 + c_1)(\sigma_x^2 + \sigma_y^2 + c_2)} \tag{32}$$

where, $\mu_x$, $\mu_y$ are the average of pixel intensity of window *x* and *y*, $\sigma_x$, $\sigma_y$, $\sigma_{xy}$ are standard deviation of window *x* and *y*, and covariance of two windows, $c_1$, and $c_2$ are $(0.01 \times L)^2$ and $(0.03 \times L)^2$, respectively, and $L$ equals to the data range of the lifetime image. Then *SSIM(x,y)* will be averaged over all the area. The average of SSIM, $\overline{\text{SSIM}}$, will be outputted, serving as "SSIM" for each pair of images in this work.

**Mean-squared errors (MSE) and peak signal-to-noise ratio (PSNR)**[9]. PSNR is a method used for measuring the quality of the image comparing to the reference image. Higher PSNR (higher score) represents better image quality. Given the reference *n x m* FLIM image *I* and the reconstructed FLIM image *K*,

$$MSE = \frac{1}{mn} \sum_{i=0}^{m-1} \sum_{j=0}^{n-1} [I(i,j) - K(i,j)]^2 \tag{33}$$

$$PSNR = 10 \log_{10}(\frac{MAX_I^2}{MSE}) \tag{34}$$

where, $MAX_I$ is the maximum possible lifetime value of the reference image *I*.

**Visual information fidelity (VIF)**[10]. VIF is a full-reference image quality model based on natural scene statistical (NSS) model and human visual system (HVS). In particular, VIF models the perceived picture quality as a communication channel, where the information of reference signal I(C,E|s) and distorted signal I(C,F|s) are first extracted by

$$I(\vec{C}^N; \vec{E}^N \middle| s^N) = \frac{1}{2} \sum_{i=1}^{N} \sum_{k=1}^{M} \log_2 \left(1 + \frac{s_i^2 \lambda_k}{\sigma_n^2}\right) \tag{35}$$

$$I(\vec{C}^N; \vec{F}^N \middle| s^N) = \frac{1}{2} \sum_{i=1}^{N} \sum_{k=1}^{M} \log_2 \left(1 + \frac{g_i^2 s_i^2 \lambda_k}{\sigma_v^2 + \sigma_n^2}\right) \tag{36}$$

where $s_i$ represents the $i^{th}$ positive scalars of the random filed (RF) in Gaussian Scale Mixture (GSM) model, $\mathbf{C}_u$ and $\lambda_k$ are the covariance and the corresponding eigenvalues of the Gaussian vector RF, $g_i$ is the $i^{th}$ deterministic scalar gain field in the distortion model as well as the variance of the stationary additive Gaussian noise RF, $\mathbf{C}_v$, and $\sigma_n^2$ represents the variance of the visual noise in the human visual system model. These parameters are estimated with maximum likelihood estimation.

The final VIF score is then defined as the ratio between two mutual information measures

$$VIF = \frac{\sum_{j \in channels} I\left(\vec{C}^{N,j}; \vec{F}^{N,j} \middle| s^{N,j}\right)}{\sum_{j \in channels} I\left(\vec{C}^{N,j}; \vec{E}^{N,j} \middle| s^{N,j}\right)} \tag{37}$$

where, $I\left(\vec{C}^{N,j}; \vec{E}^{N,j} \middle| s^{N,j}\right)$ and $I\left(\vec{C}^{N,j}; \vec{F}^{N,j} \middle| s^{N,j}\right)$ represent the information that can ideally be extracted from a particular channel, $j$, in the reference and test images respectively.

**Apparent and average lifetime for mixture of fluorophores**. We referred to technique note from ISS (http://www.iss.com/resources/pdf/technotes/FLIM_Using_Phasor_Plots.pdf) to define the apparent lifetime and average lifetime[11,12]. Given the multi-exponential decay model defined as follows,

$$I(t) = \sum_{i=1}^{n} \alpha_i e^{(-t/\tau_i)} \tag{38}$$

where, $\tau_i$ represents the lifetime, $\alpha_i$ is the amplitude of the components at t = 0, also called pre-exponential factor, and $n$ is the number of the components. Fractional contribution, $f_i$, of each decay histogram to the steady-state intensity is determined by

$$f_i = \frac{\alpha_i \tau_i}{\sum_j \alpha_j \tau_j} \tag{39}$$

Apparent lifetime is defined by,

$$\tau_{ap} = \sum_j \alpha_j \tau_j \tag{40}$$

Average lifetime is defined by,

$$\bar{\tau} = \frac{\alpha_1 \tau_1^2 + \alpha_2 \tau_2^2}{\alpha_1 \tau_1 + \alpha_2 \tau_2} = f_1 \tau_1 + f_2 \tau_2 \tag{41}$$

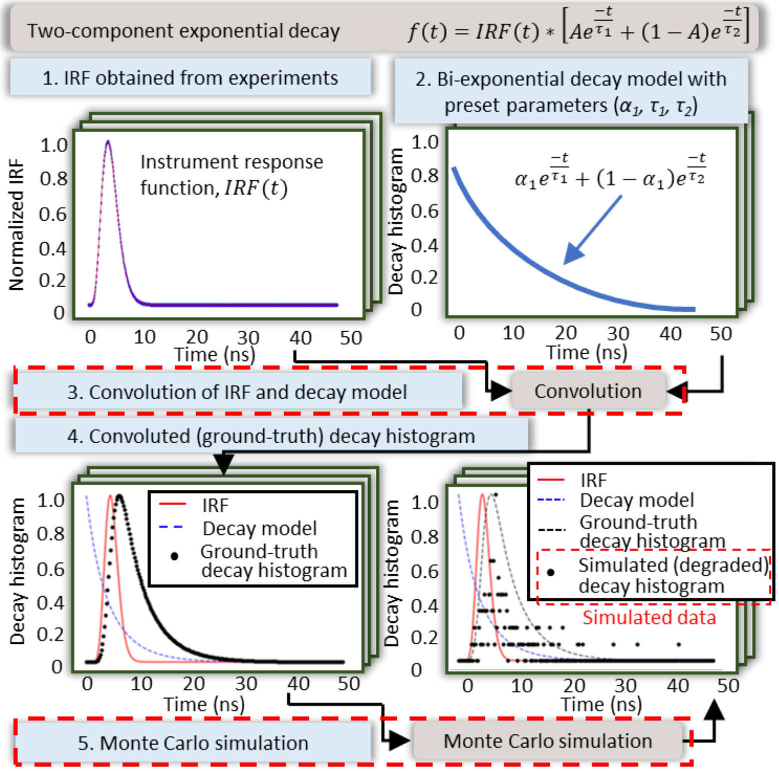


**Supplementary Fig. 1 Generation of the simulated decay histogram with lifetime parameters and photon counts by MC simulation.** First, instrument response function (IRF) was obtained from the experiments. Given the pre-defined lifetime parameters, a bi-exponential decay model is then obtained. After convolution with the experimental IRF, we normalized the convoluted decay histogram to have unit area under curve. Regarding the normalized function as probability mass function (PMF), which is served as the ground-truth decay histogram, we implemented MC simulation based on the given PMF to generate the samples with the size as the assigned photon counts. The simulated decay histogram was then obtained from the resulting samples plotted as histogram.

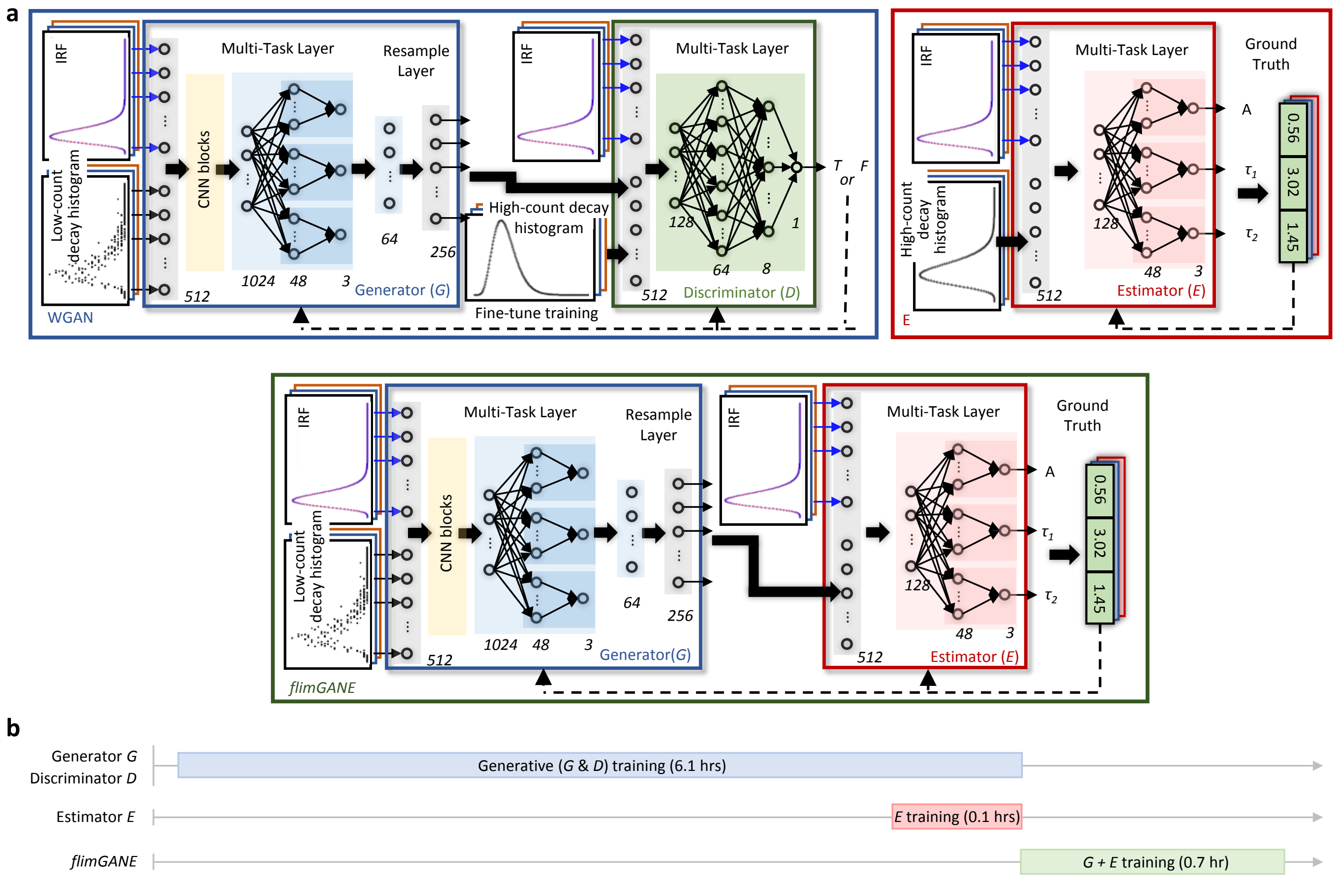


**Supplementary Fig. 2 *flimGANE*, a fluorescence lifetime analysis algorithm, can generate accurate FLIM images well-trained three subnets, generator (*G*), discriminator (*D*) and estimator (*E*).** (**a**) Schematic of *flimGANE* models. In generator (*G*), the inputs, IRF and fluorescence decay histogram, are fed into convolutional blocks followed by the multi-task layer and resample layer to reconstruct the high-photon-count fluorescence decay histogram. In discriminator (*D*), a multilayer perceptron was used to map the incoming decay histogram into a space that identified whether the input is real or fake. In estimator (*E*), a multi-task layer was employed to estimate the corresponding fluorescence lifetime parameters, fraction amplitude of shorter lifetime species ($\alpha_1$), lifetime of shorter lifetime species ($\tau_1$), and lifetime of longer lifetime species ($\tau_2$). (**b**) Timeline of *flimGANE* training process. The *G* training and *E* training processes can be performed in parallel. Then we combine the *G* and *E* together to conduct *flimGANE* training process, which is the last step before the real-world implementation.

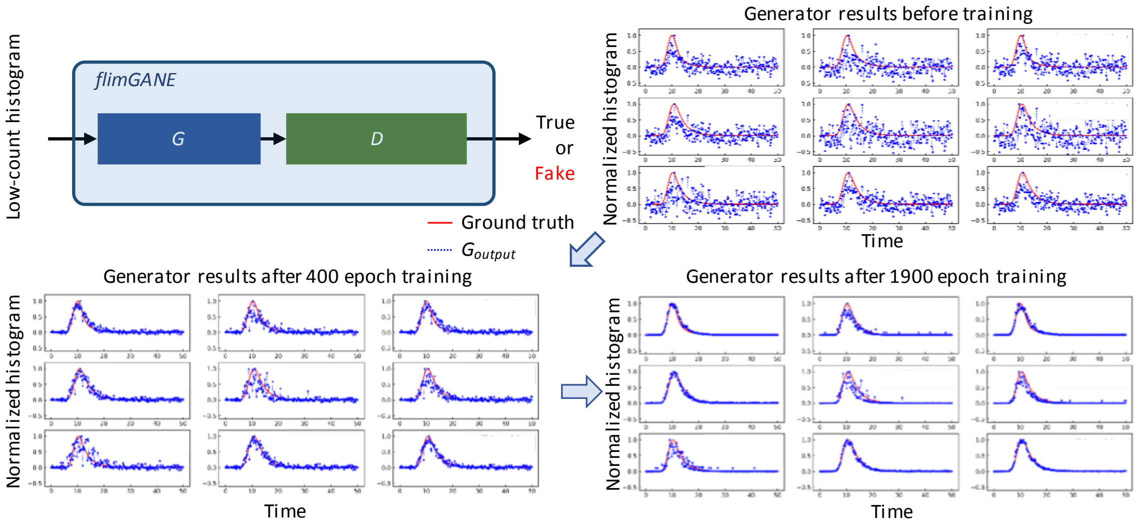


**Supplementary Fig. 3 Generator (*G*), a subnet from *flimGANE*, can transform low-photon-count decay histogram into high-photon-count one.** Given the *flimGANE* framework, the normalized low-photon-count decay histogram was transformed into the normalized ground-truth mimicking histogram. At the beginning of the training stage, the output from the generator was chaotic. The generator-inferred fluorescence decay histogram gradually matched with the ground truth.

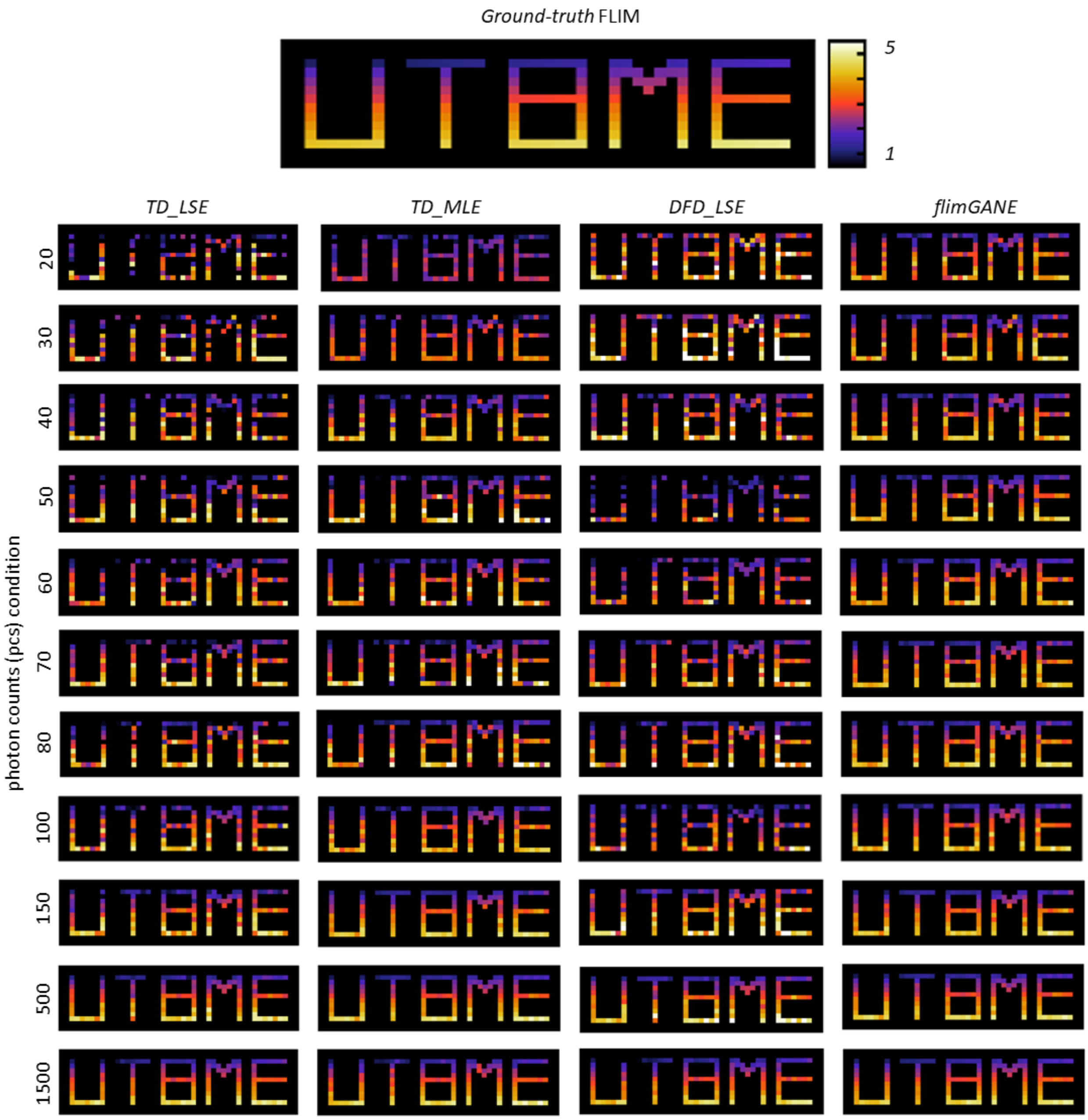


**Supplementary Fig. 4 *flimGANE* outperformed other analysis methods in FLIM images reconstruction at various photon-count conditions *in-silico.*** FLIM image was generated through MC simulation with different conditions (lifetime, $\tau$ = 1.0 ~ 5.0 ns; photon counts, *pcs* = 20, 30, 40, 50, 60, 70, 80, 100, 150, 500, 1500). It was obvious that *flimGANE* outperformed other analysis methods. As expected, under extremely low-photo-count condition (< 80 counts), *TD_MLE*, *TD_LSE*, and *DFD_LSE* are unable to make accurate estimation. Under low-photon-count condition (100, 150 counts), *TD_MLE* was close to the accurate estimation; however, *TD_LSE* and *DFD_LSE* can only generate accurate FLIM image with high-photon-count conditions (> 500 counts).

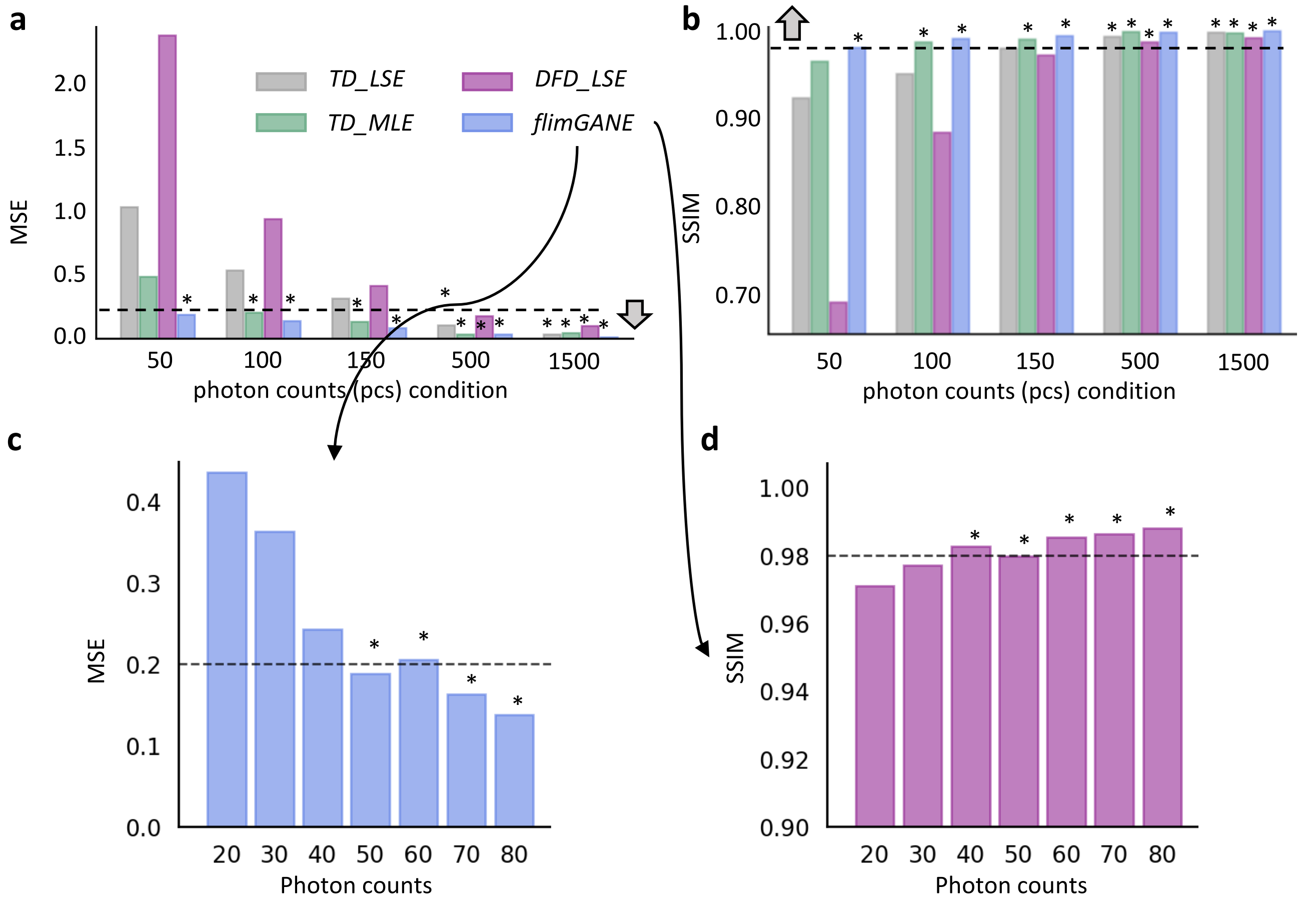


**Supplementary Fig. 5 The performance of different analysis methods was evaluated by MSE and SSIM.** The reconstructed "UT BME" FLIM images obtained by different analysis methods were evaluated by mean squared error (MSE, threshold = 0.2) and structural similarity index (SSIM, threshold = 0.98). (**a**-**b**) The performance of different analysis methods was evaluated under five conditions, 50, 100, 150, 500, 1500 photon counts. Under extremely low-photon-count condition (50 counts), the MSE and SSIM between the reconstructed FLIM images from *flimGANE* and the ground-truth FLIM images were less than 0.2 and larger than 0.98. *TD_MLE* provided MSE less than 0.2 and SSIM larger than 0.98 when the photon counts were greater than 100. *TD_LSE* and *DFD_LSE* could provide MSE less than 0.2 as the photon counts were greater than 500. (**c**-**d**) The performance boundary of *flimGANE* was further investigated. *flimGANE* could still provide MSE less than 0.2 and SSIM greater than 0.98 as the photon counts is greater than 50.

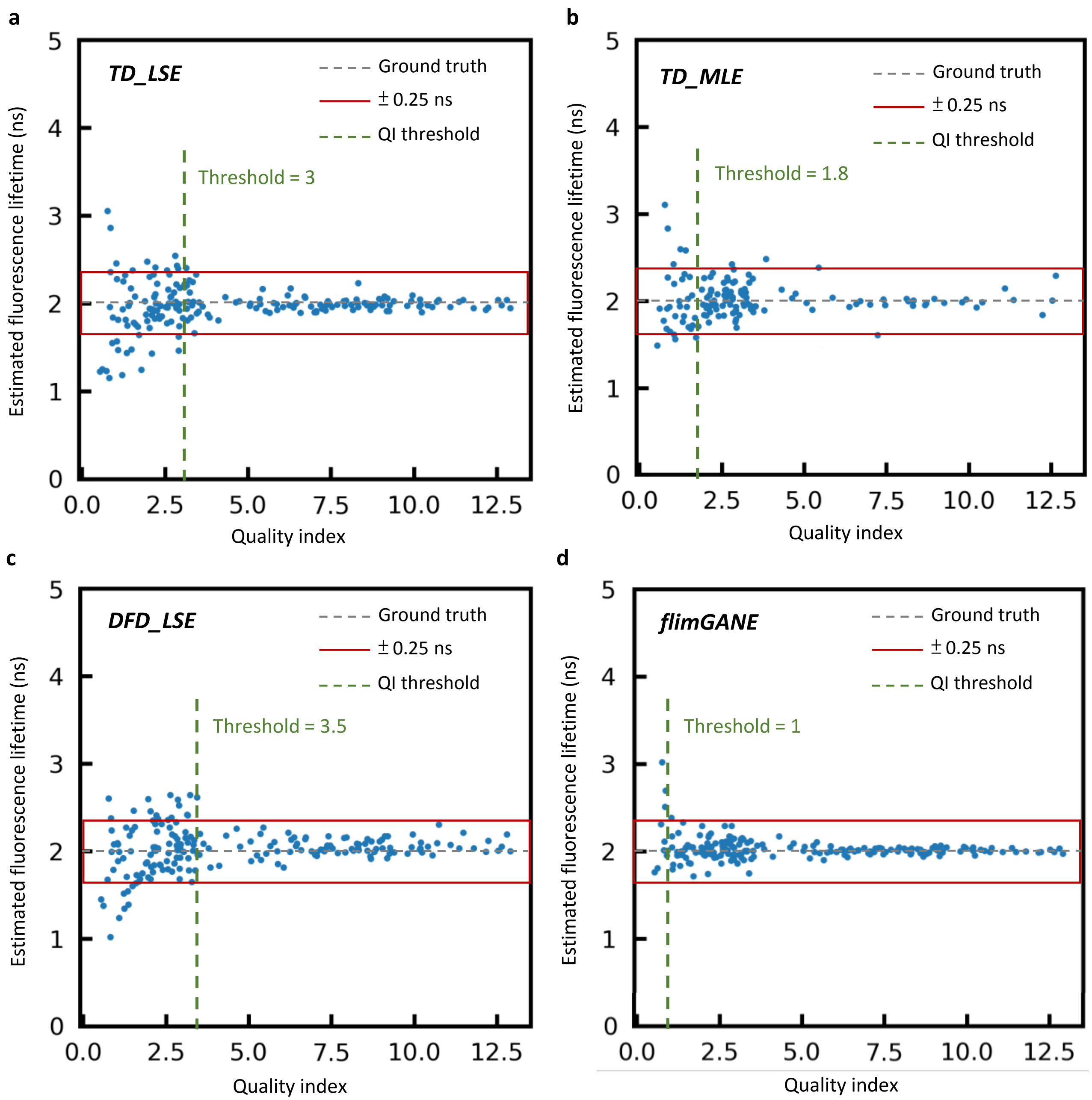


**Supplementary Fig. 6 The performance of different analysis methods was evaluated by quality index (QI).** *flimGANE* outperforms other analysis methods in fluorescence lifetime estimation from low-photon-count (noisy) decay histogram. The fluorescence decay histograms were simulated with 2 ns as pre-defined lifetime and various photon count conditions (100 ~ 10k counts), generating the simulated decay histogram with quality index (QI) ranging from 0.5 to 13.0. The tolerance of the lifetime estimate is 0.25 ns (red box). We can observe that *flimGANE* enables accurate lifetime estimation under poor QI condition as 1, which is much smaller than *TD_MLE* (1.8), *TD_LSE* (3), and *DFD_LSE* (3.5).

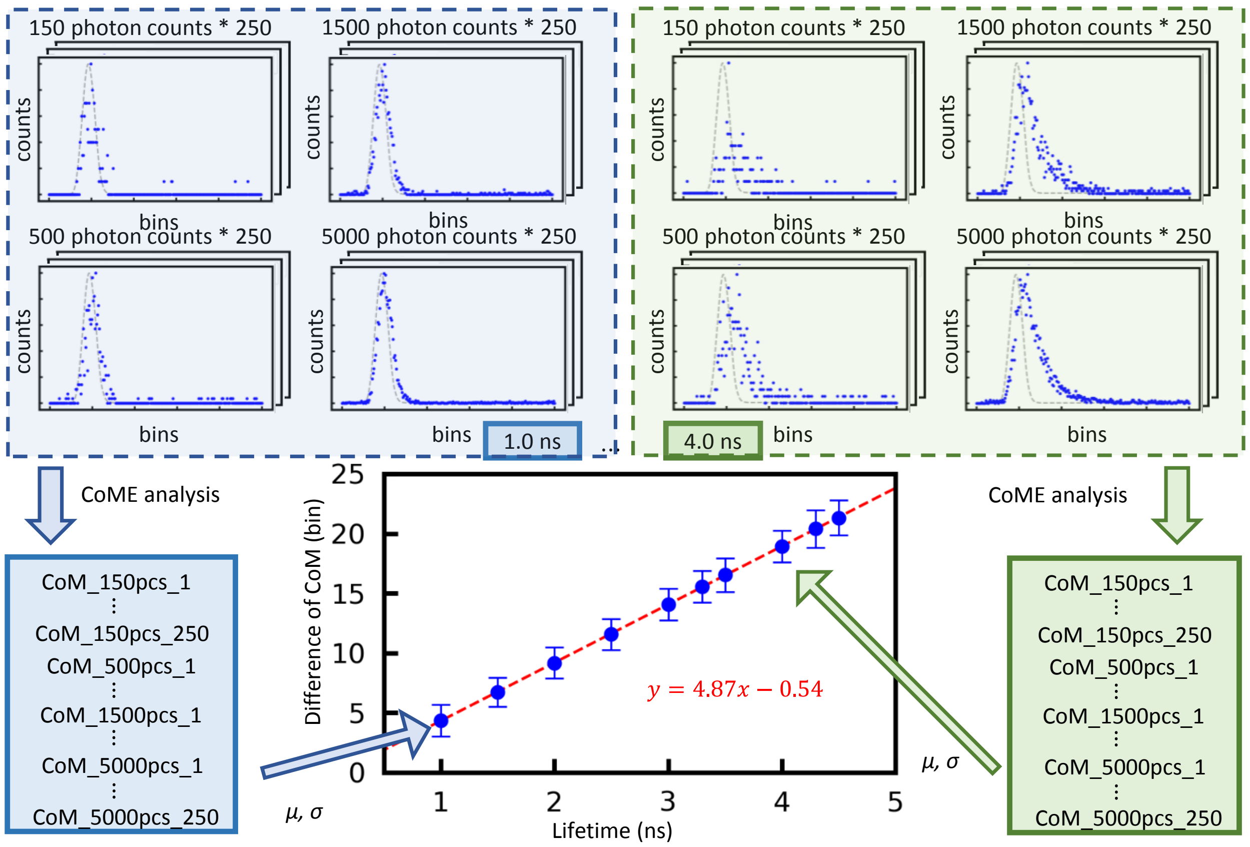


**Supplementary Fig. 7 Difference of center of mass (CoM) between the IRF and decay histogram was demonstrated positively correlated.** In order to calibrate the temporal shift between IRF and fluorescence decay histogram, we generated a calibration line to adjust the shift difference between them. First, we performed MC simulation to generate 1,000 simulated decay histograms for each lifetime value (1.0 ~ 4.5 ns) at different conditions (150, 500, 1500, 5000 photon counts). Second, we performed CoM analysis to obtain the mass center of each decay histogram and IRF. All the analysis results were plotted with the x axis as lifetime value and y axis as the distance of mass centers between fluorescence decay histogram and IRF (error bars, standard deviation, n = 1000). A linear regression line, calibration line, is obtained by fitting the analysis results.

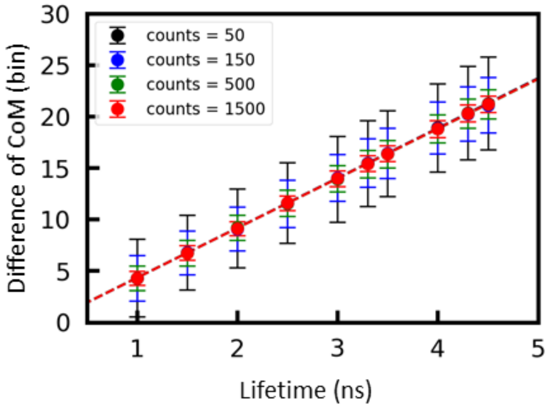


**Supplementary Fig. 8 Calibration line for center of mass evaluation (CoME) was consistent over different photon counts.** The analysis results in **Supplementary Fig. 7** were plotted with distinct photon count conditions (error bars, standard deviation, n = 250). It was demonstrated that the calibration is independent of the photon counts. The mean of the difference of CoM between IRF and decay histograms was the same at varying conditions. Moreover, it is expected to observe a higher variance at low-photon-count condition.

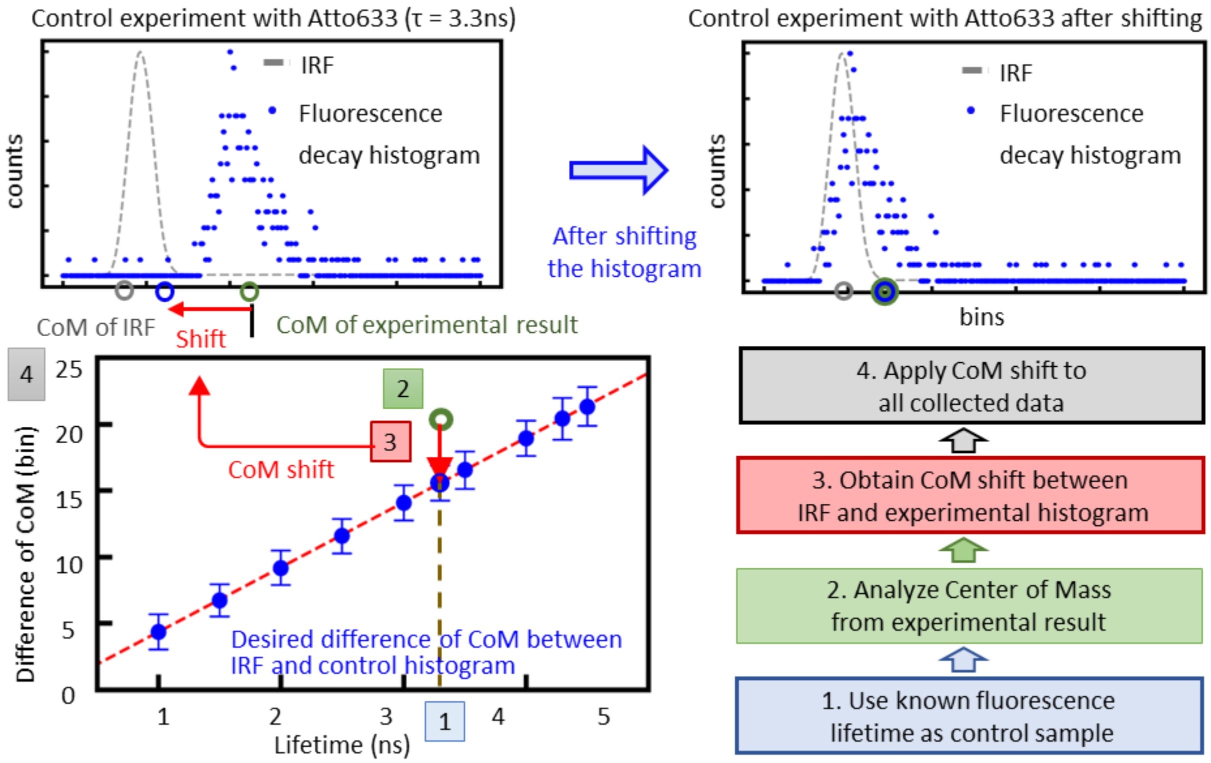


**Supplementary Fig. 9 A real-world application of the CoME.** Experimental fluorescence decay histogram aligned well with IRF based on CoME. Given the fluorescent dye with known lifetime (theoretical fluorescence lifetime of Atto633 = 3.3 ns), the desired difference of mass centers (unit: bins) of Atto633 was calculated. CoME analysis was then performed on the experimental IRF and fluorescence decay histogram to obtain the raw difference of CoM. Based on the calibration line, comparing the raw difference with the desired difference of CoM, the temporal shift can be determined with the calibration line, and the experimental fluorescence decay histogram can be reversely shifted back to the desired location. Then, shift was applied to all data collected in the same day.

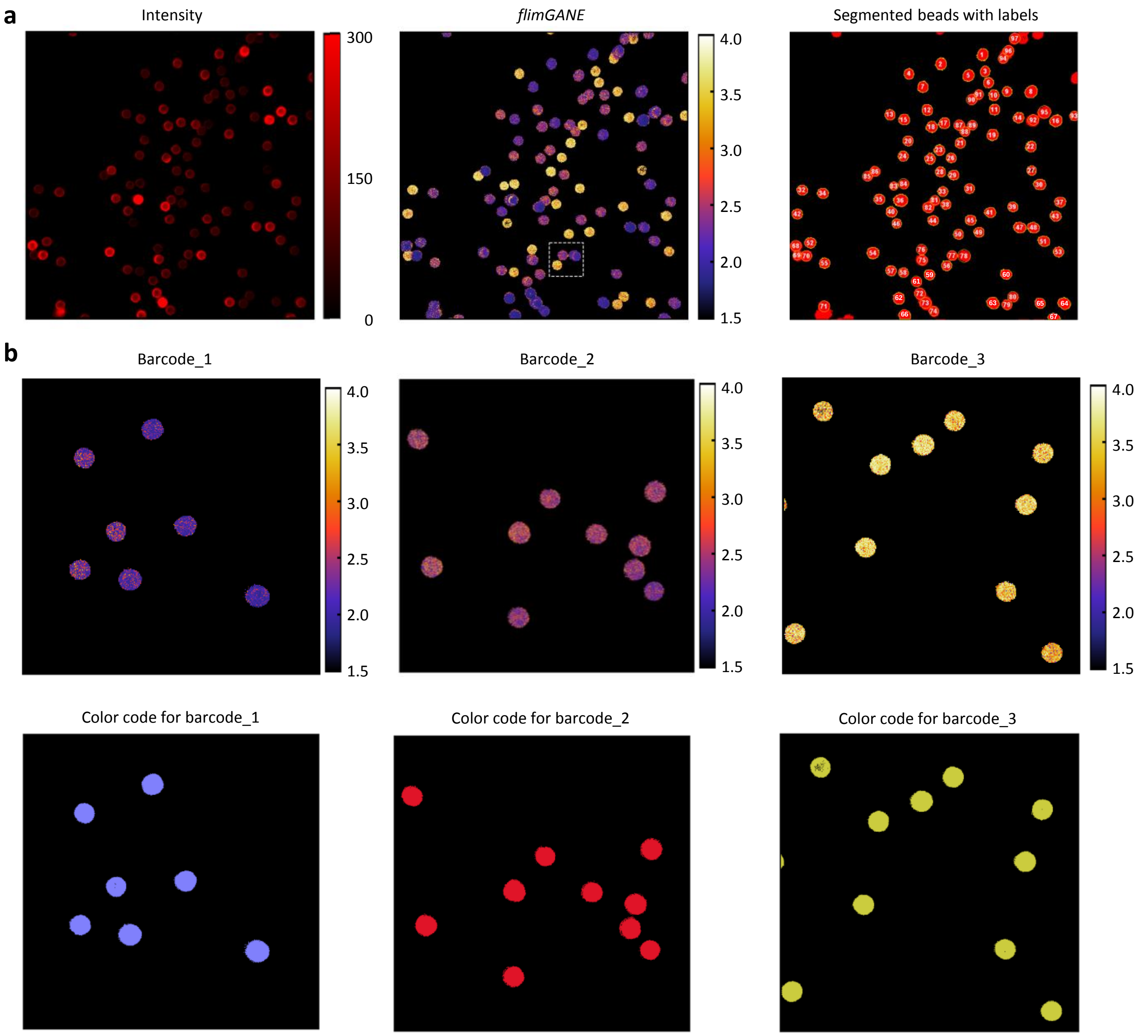


**Supplementary Fig. 10 *flimGANE* can accurately discriminate fluorescence lifetime barcode beads individually at low-photon-count condition.** (**a**) To better quantify the statistical result of FLIM barcode imaging, we selected the appropriate ROIs. Based on the intensity contrast image, we utilized ImageJ to obtain the ROIs for barcode bead. Then we manually adjusted some miss-selections and labelled each bead with a unique ID. The corresponding mean lifetime, size, and intensity information for each bead can then be extracted and recorded for further analysis. Here we can classify all beads into three different types of barcodes. (**b**) The FLIM image of each barcode was taken by ISS Alba v5, respectively, and reconstructed by *flimGANE*. The apparent lifetime of each barcode aligned well with the result obtained from *flimGANE* in (**a**).

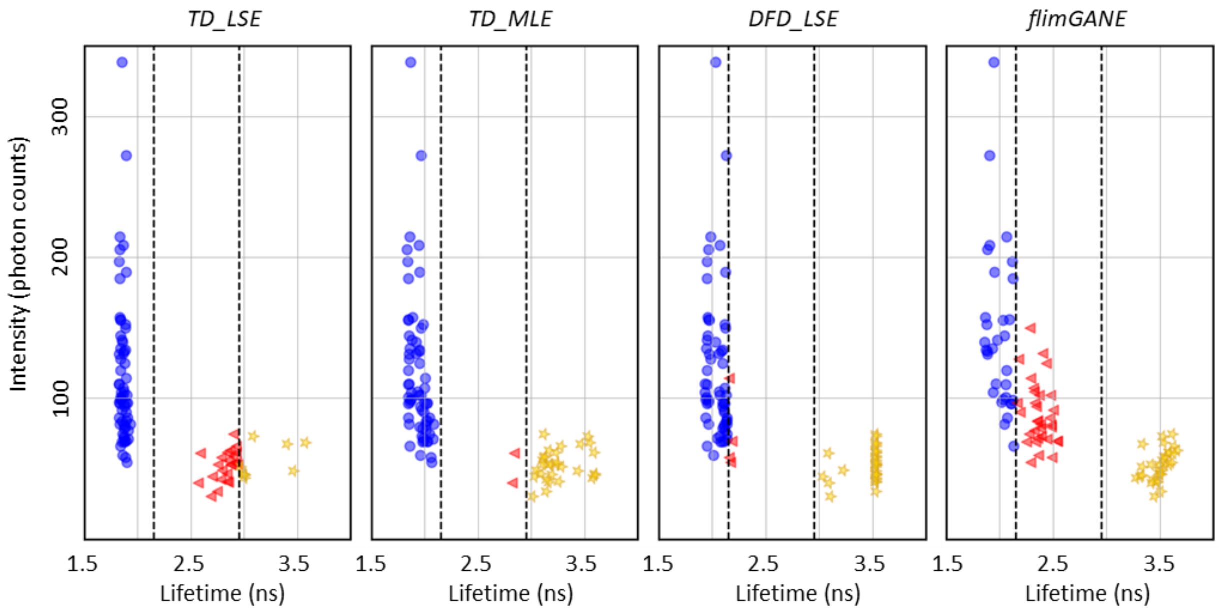


**Supplementary Fig. 11 Two dimensional (intensity versus lifetime) scattered plots from different methods illustrated the success of *flimGANE* for identifying three populations of barcodes with nearly the same size.** Each data point represents the statistical information of each bead analyzed by *TD_LSE*, *TD_MLE*, *DFD_LSE* and *flimGANE*, , respectively. Mean intensity is derived by averaging the photon counts per pixel, and the mean lifetime is obtained from the Gaussian model fitting of apparent lifetime histogram generated by different analytical methods. Based on lifetime, beads are categorized into three groups: *barcode_1* (blue) for lifetime less than 2.15 ns, *barcode_2* (red) for lifetime lying within 2.15 and 2.95 ns, and *barcode_3* (yellow) for lifetime longer than 2.95 ns.

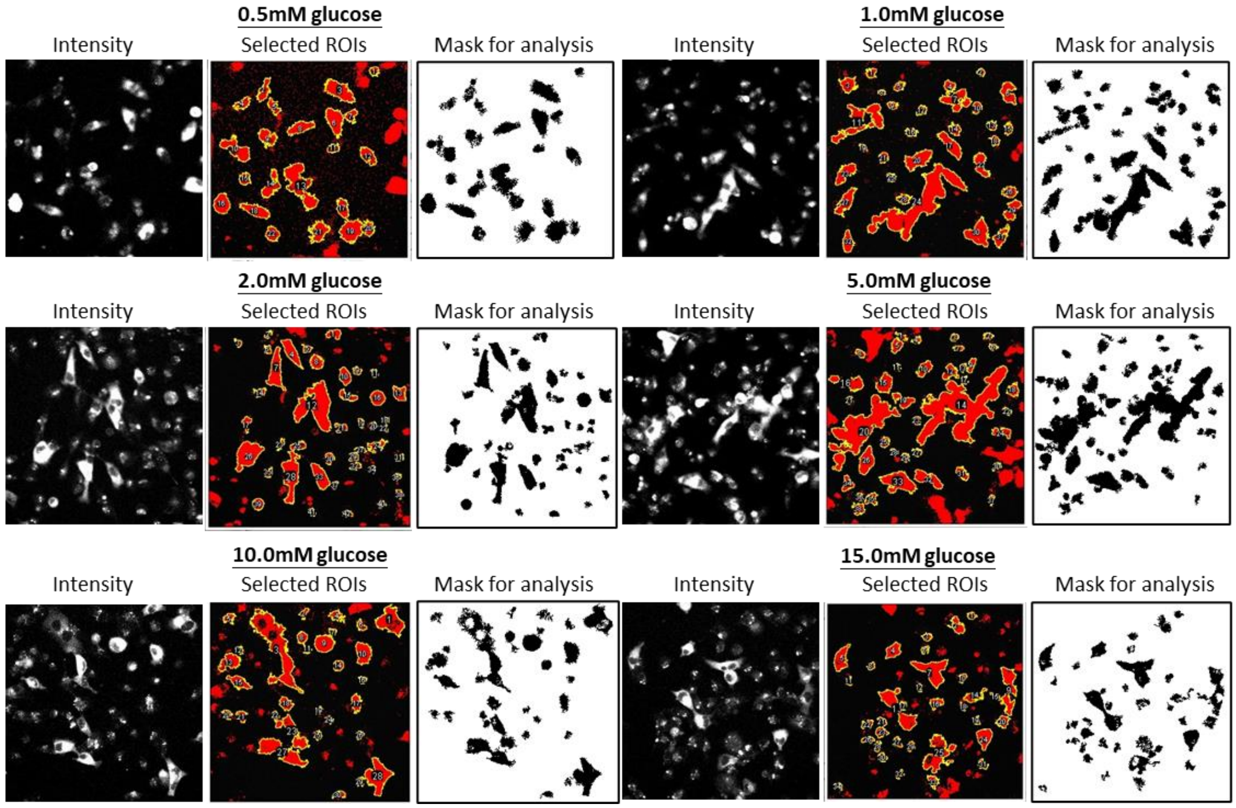


**Supplementary Fig. 12 CFP-g-YFP-transfected cells were selected from fluorescence image by ImageJ (ROI Manager).** Since the signal-to-noise ratio of FLIM images for live cell FRET experiment was low, we had to performed ROI selection to quantify the fluorescence lifetime of CFP and YFP for the cells only by ImageJ, avoiding the misinterpretation introduced by the artifact. For each experimental condition (0.5, 1.0, 2.0, 5.0, 10.0, 15.0 mM glucose), we selected ROIs independently and generated the individual mask for further lifetime analysis.

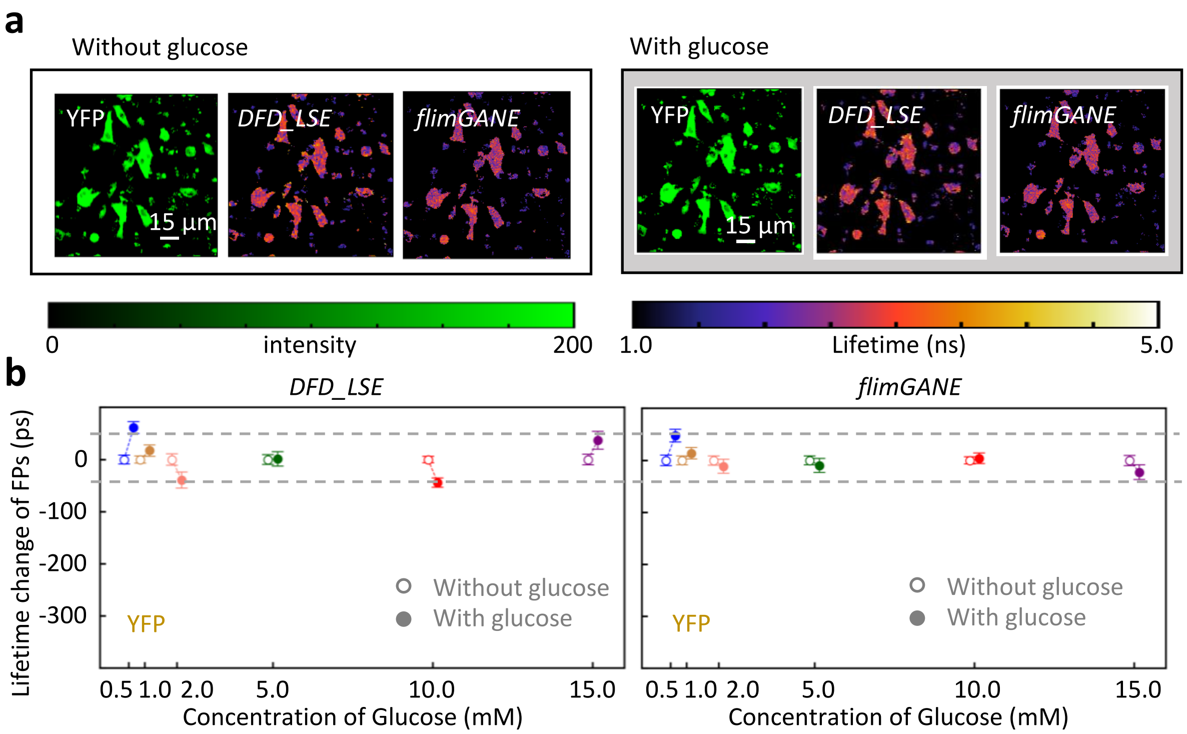


**Supplementary Fig. 13 The quantification of CFP-*g*-YFP-transfected MDA-MB-231 cell FLIM images for YFP channel**. (**a**) CFP-*g*-YFP-transfected MDA-MB-231 cells FLIM images (w/o and w/ 2 mM glucose) for YFP channel were reconstructed by *flimGANE* and *DFD_LSE*. The images of live MDA-MB-231 cells incubated in 2 mM glucose was taken with the same field of view (FOV) as the images of cells incubated in culture medium without glucose. (**b**) The mean lifetime difference between the group without and with glucose was plotted versus the six concentrations of glucose (0.5, 1.0, 2.0, 5.0, 10.0, 15.0 mM). The variation of mean lifetime difference obtained by *flimGANE* before and after adding glucose was smaller ( < ± 0.05 ns) than that obtained by *DFD_LSE*. The error bars represented the coefficient of variation (n = 2290~6824 pixels).

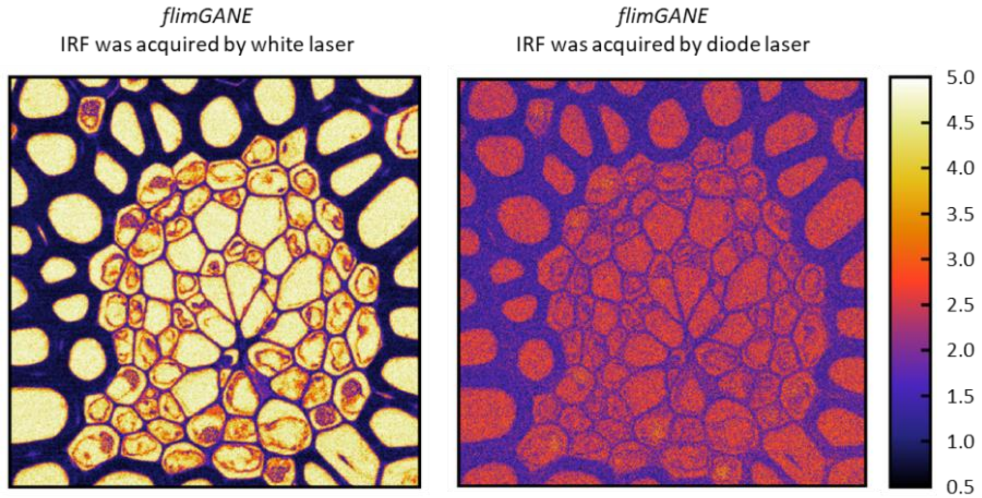


**Supplementary Fig. 14 Incorrect Convallaria FLIM image was reconstructed by *flimGANE* with incorrect IRF input.** The IRF was determined by the laser, the detector and the FLIM system electronics used in experiments. To avoid complications, traditional software tools might use a synthesized IRF to perform the analysis. However, here we want to emphasize that IRF should be considered an important factor for fluorescence lifetime analysis. The IRF used for left figure was acquired by white laser while the right one was acquired by diode laser. Although the shape visualized on the two images were similar, the estimated lifetime value for each pixel was different. It was obvious that the FLIM image reconstructed by incorrect IRF led to incorrect judgement for lifetime-based technique.

**Supplementary Table 1. Training database for different applications**

| Parameters | Fig. 1 (two dyes mixing in solution) | Fig. 2 (Barcode beads) | Fig. 3 (Convallaria) | Fig. 3 (HeLa cell) SI Fig. 17 (YFP) | Fig. 4 (CFP) | Fig. 5 (Auto-fluorescence – FAD) | Fig. 5 (Auto-fluorescence – NADH) |
|---|---|---|---|---|---|---|---|
| $\alpha_1$ | 0.0 ~ 1.0 | 0.0 ~ 1.0 | 0.98 ~ 1.00 | 0.98 ~ 1.00 | 0.0 ~ 1.0 | 0.0 ~ 1.0 | 0.0 ~ 1.0 |
| $\tau_1$ (ns) | 0.5 ~ 0.7 | 1.8 ~ 2.0 | 0.05 ~ 10.0 | 0.5 ~ 5.0 | 1.1 | 0.3 | 0.4 |
| $\tau_2$ (ns) | 3.2 ~ 3.4 | 3.4 ~ 3.6 | N/A | N/A | 3.6 | 3.0 | 2.4 ~ 4.0 |
| *IRF with excitation laser* | diode laser | diode laser | white laser | diode laser | diode laser | diode laser | diode laser |
| *rate* | 50, 100, 500, 1,500, 5,000 photons per pixel | | | | | | |
| *No. of degraded decays* | 990 k | 990 k | 300 k | 69 k | 55 k | 55 k | 467.5 k |
| *No. of ground truths* | 99 | 99 | 600 | 138 | 11 | 11 | 187 |
| *Training time for G and D (hrs)* | 18 | 18 | 6.1 | 1.5 | 1.4 | 1.4 | 6.2 |
| *Training time for E (hrs)* | 0.1 | 0.1 | 0.1 | 0.1 | 0.1 | 0.1 | 0.1 |
| *Training time for combing G and E (hrs)* | 0.8 | 0.8 | 0.7 | 0.6 | 0.1 | 0.1 | 0.7 |
| *Total model training time (hrs)* | 18.9 | 18.9 | 6.9 | 2.2 | 1.6 | 1.6 | 7.0 |

**Supplementary Table 2. Mean squared error (MSE) comparison of different analysis methods *in silico***

| Condition (photons per pixel) | ***TD_LSE*** | ***TD_MLE*** | ***DFD_LSE*** | ***flimGANE*** |
|---|---|---|---|---|
| 20 | 2.32 | 1.60 | 1.26 | 0.44 |
| 30 | 1.44 | 0.69 | 1.41 | 0.36 |
| 40 | 1.17 | 0.44 | 0.85 | 0.24 |
| 50 | 1.04 | 0.49 | 2.41 | 0.19 |
| 60 | 0.83 | 0.56 | 1.11 | 0.20 |
| 70 | 0.57 | 0.66 | 0.44 | 0.16 |
| 80 | 0.71 | 0.46 | 0.60 | 0.14 |
| 100 | 0.54 | 0.20 | 0.95 | 0.14 |
| 150 | 0.32 | 0.07 | 0.51 | 0.08 |
| 500 | 0.11 | 0.03 | 0.15 | 0.04 |
| 1500 | 0.04 | 0.05 | 0.06 | 0.01 |

**Supplementary Table 3. Peak signal-to-noise ratio (PSNR) comparison of different analysis methods *in silico***

| Condition (photons per pixel) | *TD_LSE* | *TD_MLE* | *DFD_LSE* | *flimGANE* |
|---|---|---|---|---|
| 20 | 10.32 | 11.93 | 13.57 | 17.59 |
| 30 | 12.40 | 15.59 | 14.98 | 18.38 |
| 40 | 13.30 | 17.58 | 14.18 | 20.12 |
| 50 | 13.79 | 17.06 | 10.17 | 21.23 |
| 60 | 14.80 | 16.50 | 15.86 | 20.86 |
| 70 | 16.44 | 15.77 | 13.56 | 21.86 |
| 80 | 15.47 | 17.38 | 14.39 | 22.59 |
| 100 | 16.65 | 20.87 | 14.21 | 22.51 |
| 150 | 18.94 | 22.61 | 17.73 | 24.71 |
| 500 | 23.64 | 28.72 | 21.49 | 28.54 |
| 1500 | 28.35 | 27.18 | 23.99 | 32.55 |

**Supplementary Table 4. Structural similarity index (SSIM) comparison of different analysis methods *in silico***

| Condition (photons per pixel) | *TD_LSE* | *TD_MLE* | *DFD_LSE* | *flimGANE* |
|---|---|---|---|---|
| 20 | 0.83 | 0.88 | 0.92 | 0.97 |
| 30 | 0.88 | 0.96 | 0.90 | 0.98 |
| 40 | 0.88 | 0.96 | 0.94 | 0.98 |
| 50 | 0.92 | 0.96 | 0.69 | 0.98 |
| 60 | 0.94 | 0.98 | 0.91 | 0.99 |
| 70 | 0.95 | 0.97 | 0.97 | 0.99 |
| 80 | 0.93 | 0.98 | 0.95 | 0.99 |
| 100 | 0.94 | 0.99 | 0.90 | 0.99 |
| 150 | 0.98 | 0.99 | 0.96 | 0.99 |
| 500 | 0.99 | 1.00 | 0.99 | 1.00 |
| 1500 | 1.00 | 1.00 | 0.99 | 1.00 |

**Supplementary Table 5. Goodness-of-fit reduced $\chi^2$ comparison of different analysis methods *in silico* (mean ± standard deviation)**

| Condition (photons per pixel) | *TD_LSE* | *TD_MLE* | *DFD_LSE* | *flimGANE* |
|---|---|---|---|---|
| 20 | 0.71 ±0.17 | 0.68 ±0.15 | 0.96 ±0.65 | N/A |
| 30 | 0.62 ±0.13 | 0.62 ±0.12 | 0.65 ±0.32 | N/A |
| 40 | 0.57 ±0.10 | 0.59 ±0.09 | 0.62 ±0.24 | N/A |

| | | | | |
|---|---|---|---|---|
| 50 | 0.58 ±0.11 | 0.64 ±0.10 | 1.24 ±0.48 | N/A |
| 60 | 0.61 ±0.12 | 0.68 ±0.15 | 0.50 ±0.32 | N/A |
| 70 | 0.64 ±0.13 | 0.76 ±0.18 | 0.88 ±0.36 | N/A |
| 80 | 0.69 ±0.19 | 0.83 ±0.23 | 0.68 ±0.85 | N/A |
| 100 | 0.74 ±0.17 | 0.94 ±0.26 | 0.72 ±0.38 | N/A |
| 150 | 0.90 ±0.25 | 0.98 ±0.33 | 0.50 ±0.27 | N/A |
| 500 | 0.98 ±0.18 | 1.27 ±0.27 | 0.49 ±0.53 | N/A |
| 1500 | 0.87 ±0.15 | 0.89 ±0.18 | 0.35 ±0.41 | N/A |

**Supplementary Table 6. Execution time comparison of different analysis methods *in silico***

| TD_LSE (ms / pixel) | TD_MLE (ms / pixel) | DFD_LSE (ms / pixel) | *flimGANE* (ms / pixel) |
|---|---|---|---|
| 82.40 | 906.37 | 3.94 | 0.32 |

**Supplementary Table 7. Summary of different analysis methods for the mixture of two fluorescent dyes (* mean ± standard deviation from Gaussian distributing fitting)**

| Ratio (Cy5:Atto 633) | Photons per pixel | Theoretical apparent lifetime, $\tau_\alpha$ (ns) | *TD_LSE** (ns) | *TD_MLE** (ns) | *DFD_LSE** (ns) | *flimGANE** (ns) |
|---|---|---|---|---|---|---|
| 10:0 | 178 | 0.60 | 0.59 ±0.05 | 0.71 ±0.03 | 0.62 ±0.02 | 0.68 ±0.15 |
| 9:1 | 135 | 0.87 | 0.81 ±0.14 | 1.11 ±0.17 | 0.70 ±0.06 | 0.88 ±0.10 |
| 8:2 | 127 | 1.14 | 1.05 ±0.19 | 1.28 ±0.20 | 0.80 ±0.09 | 1.13 ±0.11 |
| 7:3 | 121 | 1.41 | 1.02 ±0.19 | 1.52 ±0.23 | 0.95 ±0.13 | 1.42 ±0.14 |
| 6:4 | 127 | 1.68 | 1.10 ±0.22 | 1.82 ±0.27 | 1.09 ±0.21 | 1.70 ±0.17 |
| 5:5 | 120 | 1.95 | 1.53 ±0.26 | 1.92 ±0.26 | 1.21 ±0.28 | 1.99 ±0.25 |
| 4:6 | 129 | 2.22 | 1.68 ±0.27 | 2.32 ±0.33 | 1.29 ±0.29 | 2.26 ±0.25 |
| 3:7 | 134 | 2.49 | 1.74 ±0.28 | 2.49 ±0.34 | 1.99 ±1.20 | 2.64 ±0.37 |
| 2:8 | 137 | 2.76 | 1.80 ±0.33 | 2.85 ±0.37 | 2.14 ±1.27 | 2.82 ±0.20 |
| 0:10 | 190 | 3.30 | 2.67 ±0.60 | 3.32 ±0.06 | 3.04 ±1.05 | 3.31 ±0.30 |

**Supplementary Table 8. Summary of each detected bead**

| ID | *TD_LSE* $\tau_\alpha$ (ns) | *TD_LSE* (Barcode) | *TD_MLE* $\tau_\alpha$ (ns) | *TD_MLE* (Barcode) | *DFD_LSE* $\tau_\alpha$ (ns) | *DFD_LSE* (Barcode) | *flimGANE* $\tau_\alpha$ (ns) | *flimGANE* (Barcode) |
|---|---|---|---|---|---|---|---|---|
| 1 | 1.83 | 1 | 1.85 | 1 | 1.93 | 1 | 1.96 | 1 |
| 2 | 1.83 | 1 | 1.86 | 1 | 1.97 | 1 | 2.02 | 1 |
| 3 | 2.84 | 2 | 3.26 | 3 | 3.52 | 3 | 3.47 | 3 |
| 4 | 2.85 | 2 | 3.11 | 3 | 3.52 | 3 | 3.52 | 3 |
| 5 | 1.87 | 1 | 2.00 | 1 | 2.12 | 1 | 2.31 | 2 |

| | | | | | | | | |
|---|---|---|---|---|---|---|---|---|
| 6 | 2.94 | 2 | 3.23 | 3 | 3.52 | 3 | 3.54 | 3 |
| 7 | 1.83 | 1 | 1.85 | 1 | 1.96 | 1 | 2.05 | 1 |
| 8 | 1.84 | 1 | 1.84 | 1 | 1.97 | 1 | 2.09 | 1 |
| 9 | 2.85 | 2 | 3.16 | 3 | 3.52 | 3 | 3.48 | 3 |
| 10 | 1.85 | 1 | 1.98 | 1 | 2.10 | 1 | 2.33 | 2 |
| 11 | 1.87 | 1 | 1.97 | 1 | 2.07 | 1 | 2.38 | 2 |
| 12 | 1.90 | 1 | 1.97 | 1 | 2.13 | 1 | 2.47 | 2 |
| 13 | 1.88 | 1 | 1.99 | 1 | 2.19 | 2 | 2.53 | 2 |
| 14 | 2.89 | 2 | 3.13 | 3 | 3.52 | 3 | 3.54 | 3 |
| 15 | 1.83 | 1 | 1.84 | 1 | 1.94 | 1 | 2.07 | 1 |
| 16 | 1.88 | 1 | 1.95 | 1 | 2.10 | 1 | 2.43 | 2 |
| 17 | 1.84 | 1 | 1.84 | 1 | 1.94 | 1 | 2.05 | 1 |
| 18 | 1.87 | 1 | 1.93 | 1 | 2.06 | 1 | 2.32 | 2 |
| 19 | 1.83 | 1 | 1.84 | 1 | 1.95 | 1 | 2.06 | 1 |
| 20 | 1.84 | 1 | 1.86 | 1 | 1.95 | 1 | 2.13 | 1 |
| 21 | 2.90 | 2 | 3.11 | 3 | 3.52 | 3 | 3.61 | 3 |
| 22 | 1.87 | 1 | 1.98 | 1 | 2.10 | 1 | 2.38 | 2 |
| 23 | 1.85 | 1 | 1.97 | 1 | 2.11 | 1 | 2.50 | 2 |
| 24 | 1.88 | 1 | 1.95 | 1 | 2.11 | 1 | 2.48 | 2 |
| 25 | 1.87 | 1 | 1.92 | 1 | 2.06 | 1 | 2.40 | 2 |
| 26 | 2.87 | 2 | 3.18 | 3 | 3.52 | 3 | 3.58 | 3 |
| 27 | 2.68 | 2 | 3.01 | 3 | 3.10 | 3 | 3.45 | 3 |
| 28 | 2.93 | 2 | 3.28 | 3 | 3.52 | 3 | 3.63 | 3 |
| 29 | 1.87 | 1 | 2.01 | 1 | 2.11 | 1 | 2.48 | 2 |
| 30 | 1.84 | 1 | 1.85 | 1 | 1.95 | 1 | 2.06 | 1 |
| 31 | 2.78 | 2 | 3.12 | 3 | 3.52 | 3 | 3.54 | 3 |
| 32 | 2.84 | 2 | 3.23 | 3 | 3.52 | 3 | 3.58 | 3 |
| 33 | 1.89 | 1 | 2.05 | 1 | 2.15 | 2 | 2.49 | 2 |
| 34 | 1.84 | 1 | 1.87 | 1 | 1.96 | 1 | 2.12 | 1 |
| 35 | 1.84 | 1 | 1.85 | 1 | 1.99 | 1 | 2.17 | 2 |
| 36 | 1.85 | 1 | 1.86 | 1 | 2.03 | 1 | 1.94 | 1 |
| 37 | 1.86 | 1 | 2.04 | 1 | 2.13 | 1 | 2.28 | 2 |
| 38 | 2.84 | 2 | 3.09 | 3 | 3.52 | 3 | 3.50 | 3 |
| 39 | 2.99 | 3 | 3.59 | 3 | 3.52 | 3 | 3.49 | 3 |
| 40 | 1.83 | 1 | 1.85 | 1 | 1.95 | 1 | 2.10 | 1 |

| | | | | | | | | |
|---|---|---|---|---|---|---|---|---|
| 41 | 1.89 | 1 | 2.02 | 1 | 2.14 | 1 | 2.35 | 2 |
| 42 | 2.93 | 2 | 3.59 | 3 | 3.52 | 3 | 3.58 | 3 |
| 43 | 3.08 | 3 | 3.53 | 3 | 3.52 | 3 | 3.52 | 3 |
| 44 | 1.91 | 1 | 2.00 | 1 | 2.11 | 1 | 2.33 | 2 |
| 45 | 1.88 | 1 | 1.96 | 1 | 2.10 | 1 | 2.34 | 2 |
| 46 | 2.75 | 2 | 3.13 | 3 | 3.52 | 3 | 3.51 | 3 |
| 47 | 1.87 | 1 | 1.95 | 1 | 2.08 | 1 | 1.88 | 1 |
| 48 | 1.84 | 1 | 1.85 | 1 | 1.96 | 1 | 2.04 | 1 |
| 49 | 3.57 | 3 | 3.55 | 3 | 3.52 | 3 | 3.54 | 3 |
| 50 | 2.88 | 2 | 3.17 | 3 | 3.52 | 3 | 3.49 | 3 |
| 51 | 1.89 | 1 | 1.99 | 1 | 2.11 | 1 | 2.19 | 2 |
| 52 | 1.84 | 1 | 1.87 | 1 | 1.94 | 1 | 2.03 | 1 |
| 53 | 2.70 | 2 | 3.03 | 3 | 3.03 | 3 | 3.31 | 3 |
| 54 | 1.83 | 1 | 1.84 | 1 | 1.95 | 1 | 2.12 | 1 |
| 55 | 1.90 | 1 | 2.05 | 1 | 2.14 | 1 | 2.47 | 2 |
| 56 | 2.75 | 2 | 3.11 | 3 | 3.52 | 3 | 3.42 | 3 |
| 57 | 1.85 | 1 | 2.01 | 1 | 2.13 | 1 | 2.35 | 2 |
| 58 | 2.78 | 2 | 3.15 | 3 | 3.52 | 3 | 3.43 | 3 |
| 59 | 1.93 | 1 | 2.08 | 1 | 2.13 | 1 | 2.38 | 2 |
| 60 | 1.87 | 1 | 2.02 | 1 | 2.08 | 1 | 2.25 | 2 |
| 61 | 1.91 | 1 | 2.08 | 1 | 2.13 | 1 | 2.44 | 2 |
| 62 | 1.88 | 1 | 1.95 | 1 | 2.09 | 1 | 1.87 | 1 |
| 63 | 1.84 | 1 | 1.86 | 1 | 1.95 | 1 | 1.93 | 1 |
| 64 | 1.89 | 1 | 2.02 | 1 | 2.10 | 1 | 2.17 | 2 |
| 65 | 3.01 | 3 | 3.57 | 3 | 3.52 | 3 | 3.27 | 3 |
| 66 | 1.88 | 1 | 1.98 | 1 | 2.11 | 1 | 1.88 | 1 |
| 67 | 1.87 | 1 | 2.02 | 1 | 2.12 | 1 | 2.11 | 1 |
| 68 | 1.87 | 1 | 1.94 | 1 | 2.07 | 1 | 1.90 | 1 |
| 69 | 3.46 | 3 | 3.43 | 3 | 3.52 | 3 | 3.49 | 3 |
| 70 | 1.84 | 1 | 1.88 | 1 | 1.95 | 1 | 1.87 | 1 |
| 71 | 1.85 | 1 | 1.88 | 1 | 1.95 | 1 | 1.98 | 1 |
| 72 | 1.86 | 1 | 1.92 | 1 | 2.03 | 1 | 1.86 | 1 |
| 73 | 1.89 | 1 | 1.96 | 1 | 2.13 | 1 | 1.90 | 1 |
| 74 | 1.85 | 1 | 1.87 | 1 | 1.93 | 1 | 1.94 | 1 |
| 75 | 1.88 | 1 | 1.97 | 1 | 2.12 | 1 | 2.28 | 2 |

| | | | | | | | | |
|---|---|---|---|---|---|---|---|---|
| 76 | 2.79 | 2 | 3.04 | 3 | 3.22 | 3 | 3.51 | 3 |
| 77 | 1.91 | 1 | 2.01 | 1 | 2.12 | 1 | 2.32 | 2 |
| 78 | 1.84 | 1 | 1.85 | 1 | 1.97 | 1 | 2.03 | 1 |
| 79 | 2.96 | 3 | 3.28 | 3 | 3.52 | 3 | 3.33 | 3 |
| 80 | 3.01 | 3 | 3.60 | 3 | 3.52 | 3 | 3.33 | 3 |
| 81 | 1.89 | 1 | 2.00 | 1 | 2.16 | 2 | 2.29 | 2 |
| 82 | 1.89 | 1 | 1.95 | 1 | 2.12 | 1 | 1.95 | 1 |
| 83 | 2.87 | 2 | 3.17 | 3 | 3.52 | 3 | 3.66 | 3 |
| 84 | 2.92 | 2 | 3.24 | 3 | 3.52 | 3 | 3.61 | 3 |
| 85 | 2.59 | 2 | 2.84 | 2 | 3.08 | 3 | 3.63 | 3 |
| 86 | 1.83 | 1 | 1.85 | 1 | 1.97 | 1 | 1.88 | 1 |
| 87 | 1.87 | 1 | 1.95 | 1 | 2.07 | 1 | 2.37 | 2 |
| 88 | 1.89 | 1 | 2.02 | 1 | 2.12 | 1 | 2.54 | 2 |
| 89 | 2.57 | 2 | 2.82 | 2 | 3.09 | 3 | 3.42 | 3 |
| 90 | 1.87 | 1 | 1.98 | 1 | 2.04 | 1 | 2.35 | 2 |
| 91 | 1.86 | 1 | 1.96 | 1 | 2.01 | 1 | 2.36 | 2 |
| 92 | 1.83 | 1 | 1.83 | 1 | 1.96 | 1 | 1.88 | 1 |
| 93 | 3.00 | 3 | 3.19 | 3 | 3.52 | 3 | 3.51 | 3 |
| 94 | 3.40 | 3 | 3.46 | 3 | 3.53 | 3 | 3.34 | 3 |
| 95 | 1.83 | 1 | 1.84 | 1 | 1.95 | 1 | 2.12 | 1 |
| 96 | 1.83 | 1 | 1.86 | 1 | 1.98 | 1 | 2.06 | 1 |
| 97 | 1.90 | 1 | 2.06 | 1 | 2.17 | 2 | 2.29 | 2 |

**Supplementary Table 9. MSE, PSNR, SSIM, and VIF comparison of different analysis methods for *Convallaria* FLIM images (Standard: medium-count *TD_MLE* FLIM)**

| Condition (photons per pixel) | Method | MSE | PSNR | SSIM | VIF |
|---|---|---|---|---|---|
| 50 – 200 photon counts per pixel | *TD_LSE* | 2.01 | 10.94 | 0.71 | 0.17 |
| | *TD_MLE* | 0.52 | 16.84 | 0.89 | 0.22 |
| | *DFD_LSE* | 4.10 | 7.85 | 0.51 | 0.09 |
| | *flimGANE* | 0.61 | 15.71 | 0.88 | 0.22 |

**Supplementary Table 10. MSE, PSNR, SSIM and VIF comparison of different analysis methods for live HeLa cell FLIM images (Standard: medium-high-count FLIM)**

| Condition (average photons per pixel) | Method | MSE | PSNR | SSIM | VIF |
|---|---|---|---|---|---|

| | | | | | |
|---|---|---|---|---|---|
| ~180 (Red channel) | *TD_LSE* | 0.28 | 19.44 | 0.67 | 0.06 |
| | *TD_MLE* | 0.10 | 23.90 | 0.75 | 0.14 |
| | *DFD_LSE* | 0.24 | 20.26 | 0.68 | 0.11 |
| | *flimGANE* | 0.13 | 22.88 | 0.87 | 0.22 |
| ~180 (Blue channel) | *TD_LSE* | 0.37 | 18.34 | 0.93 | 0.25 |
| | *TD_MLE* | 0.12 | 23.10 | 0.95 | 0.45 |
| | *DFD_LSE* | 0.79 | 15.01 | 0.93 | 0.31 |
| | *flimGANE* | 0.10 | 24.01 | 0.98 | 0.47 |

**Supplementary Table 11. MSE, PSNR and SSIM comparison of different analysis methods for FRET FLIM images (Reference: *TD_MLE* FLIM)**

| FP type | [Glucose] (mM) | With or without glucose | Method | MSE | PSNR | SSIM |
|---|---|---|---|---|---|---|
| CFP | 0.5 | Before | *TD_LSE* | 0.69 | 15.59 | 0.98 |
| | | | *TD_MLE* | N/A | | |
| | | | *DFD_LSE* | 0.95 | 14.20 | 0.98 |
| | | | *flimGANE* | 0.29 | 19.40 | 0.99 |
| | | After | *TD_LSE* | 0.66 | 15.77 | 0.98 |
| | | | *TD_MLE* | N/A | | |
| | | | *DFD_LSE* | 0.90 | 14.43 | 0.98 |
| | | | *flimGANE* | 0.26 | 19.76 | 0.99 |
| | 1.0 | Before | *TD_LSE* | 0.61 | 16.13 | 0.98 |
| | | | *TD_MLE* | N/A | | |
| | | | *DFD_LSE* | 0.90 | 14.46 | 0.97 |
| | | | *flimGANE* | 0.27 | 19.68 | 0.99 |
| | | After | *TD_LSE* | 0.56 | 16.48 | 0.98 |
| | | | *TD_MLE* | N/A | | |
| | | | *DFD_LSE* | 0.84 | 14.75 | 0.97 |
| | | | *flimGANE* | 0.25 | 20.09 | 0.99 |
| | 2.0 | Before | *TD_LSE* | 0.57 | 16.41 | 0.97 |
| | | | *TD_MLE* | N/A | | |
| | | | *DFD_LSE* | 0.89 | 14.49 | 0.96 |
| | | | *flimGANE* | 0.26 | 19.79 | 0.98 |
| | | After | *TD_LSE* | 0.51 | 16.94 | 0.97 |
| | | | *TD_MLE* | N/A | | |
| | | | *DFD_LSE* | 0.80 | 14.94 | 0.96 |

| | | | | | | |
|---|---|---|---|---|---|---|
| | | | *flimGANE* | 0.18 | 21.36 | 0.99 |
| | 5.0 | Before | *TD_LSE* | 0.55 | 16.58 | 0.94 |
| | | | *TD_MLE* | N/A | | |
| | | | *DFD_LSE* | 0.32 | 18.93 | 0.95 |
| | | | *flimGANE* | 0.21 | 20.78 | 0.97 |
| | | After | *TD_LSE* | 0.48 | 17.19 | 0.95 |
| | | | *TD_MLE* | N/A | | |
| | | | *DFD_LSE* | 0.75 | 15.21 | 0.93 |
| | | | *flimGANE* | 0.14 | 22.57 | 0.98 |
| | 10.0 | Before | *TD_LSE* | 0.45 | 17.43 | 0.98 |
| | | | *TD_MLE* | N/A | | |
| | | | *DFD_LSE* | 0.85 | 14.67 | 0.98 |
| | | | *flimGANE* | 0.25 | 20.07 | 0.99 |
| | | After | *TD_LSE* | 0.43 | 17.61 | 0.99 |
| | | | *TD_MLE* | N/A | | |
| | | | *DFD_LSE* | 0.85 | 14.68 | 0.98 |
| | | | *flimGANE* | 0.22 | 20.51 | 0.99 |
| | 15.0 | Before | *TD_LSE* | 0.53 | 16.76 | 0.97 |
| | | | *TD_MLE* | N/A | | |
| | | | *DFD_LSE* | 0.84 | 14.73 | 0.96 |
| | | | *flimGANE* | 0.25 | 20.01 | 0.99 |
| | | After | *TD_LSE* | 0.49 | 17.06 | 0.98 |
| | | | *TD_MLE* | N/A | | |
| | | | *DFD_LSE* | 0.81 | 14.87 | 0.97 |
| | | | *flimGANE* | 0.24 | 20.11 | 0.99 |

**Supplementary Table 12. Apparent lifetime of CFP calculated with Gaussian distribution fitting (* mean ± standard deviation errors on the parameter estimate)**

| FP type | [Glucose] (mM) | With or without glucose | *TD_LSE** | *TD_MLE** | *DFD_LSE** | *flimGANE ** |
|---|---|---|---|---|---|---|
| CFP | 0.5 | Before | 1.62 ± 0.02 | 2.08 ± 0.01 | N/A | 2.11 ± 0.01 |
| | | After | 1.58 ± 0.02 | 2.08 ± 0.01 | N/A | 2.07 ± 0.01 |
| | 1.0 | Before | 1.63 ± 0.03 | 2.12 ± 0.01 | N/A | 2.22 ± 0.01 |
| | | After | 1.59 ± 0.02 | 2.06 ± 0.01 | N/A | 2.13 ± 0.01 |
| | 2.0 | Before | 1.72 ± 0.02 | 2.19 ± 0.01 | N/A | 2.25 ± 0.01 |
| | | After | 1.54 ± 0.02 | 2.05 ± 0.01 | N/A | 2.13 ± 0.01 |

| | | | | | | |
|---|---|---|---|---|---|---|
| | 5.0 | Before | 1.62 ±0.02 | 2.18 ±0.01 | N/A | 2.25 ±0.00 |
| | | After | 1.46 ±0.02 | 2.02 ±0.01 | N/A | 2.14 ±0.01 |
| | 10.0 | Before | 1.64 ±0.04 | 2.01 ±0.01 | N/A | 2.17 ±0.01 |
| | | After | 1.55 ±0.04 | 1.95 ±0.01 | N/A | 2.06 ±0.01 |
| | 15.0 | Before | 1.62 ±0.03 | 2.07 ±0.01 | N/A | 2.17 ±0.01 |
| | | After | 1.55 ±0.02 | 2.01 ±0.01 | N/A | 2.02 ±0.01 |

**Supplementary Table 13. MSE, PSNR, and SSIM comparison of different methods for autofluorescence FLIM images** (Reference: *TD_MLE* FLIM)

| Condition | Method | MSE | PSNR | SSIM |
|---|---|---|---|---|
| FAD | *TD_LSE* | 0.22 | 20.48 | 0.88 |
| | *DFD_LSE* | 0.69 | 15.59 | 0.71 |
| | *flimGANE* | 0.24 | 20.08 | 0.86 |
| NADH | *TD_LSE* | 0.66 | 15.76 | 0.66 |
| | *DFD_LSE* | 1.39 | 12.55 | 0.57 |
| | *flimGANE* | 0.21 | 20.65 | 0.87 |
